# Modelling TGFβR and Hh pathway regulation of prognostic matrisome molecules in ovarian cancer

**DOI:** 10.1101/2021.02.01.428374

**Authors:** Robin M. Delaine-Smith, Eleni Maniati, Beatrice Malacrida, Sam Nichols, Reza Roozitalab, Roanne R. Jones, Laura Lecker, Oliver M.T. Pearce, Martin M. Knight, Frances R. Balkwill

## Abstract

In a multi-level ‘deconstruction’ of omental metastases, we previously identified a prognostic matrisome gene expression signature in high-grade serous ovarian cancer (HGSOC) and twelve other malignancies. Here, our aim was to understand how six of these extracellular matrix, ECM, molecules, COL11A1, COMP, FN1, VCAN, CTSB and COL1A1, are up-regulated in cancer. Using biopsies, we identified significant associations between TGFβR activity, Hedgehog signalling and these ECM molecules and then studied the associations in mono-, co- and tri-culture. Activated omental fibroblasts produced more matrix than malignant cells, directed by TGFβR and Hedgehog signalling crosstalk. We ‘reconstructed’ omental metastases in tri-culture of HGSOC cells, omental fibroblasts and adipocytes. This combination was sufficient to generate all six ECM proteins and the matrisome expression signature. TGFβR and Hedgehog inhibitor combinations attenuated fibroblast activation, gel remodelling and ECM remodelling in these models. The tri-culture model reproduces key features of omental metastases and allows study of diseased-associated ECM.

## Introduction

Desmoplasia and extracellular matrix (ECM) remodelling are common features of human solid tumors and are driven by the continued presence of malignant cells. In high-grade serous ovarian cancer (HGSOC) there is increasing evidence that stromal components play a key role in tumor growth, promoting aggressive malignant cell phenotypes ^1,2^.

Previously, we reported a tumor-associated matrisome gene signature, the Matrix Index (MI), which predicted poor prognosis in HGSOC patients and twelve other solid tumor types ^3^. Six of these genes, *COL11A1, COMP, FN1, VCAN, CTSB* and *COL1A1*, were significantly upregulated with disease and have all previously been associated with tumor progression, poor prognosis, and invasive malignant cell phenotypes in ovarian and/or other cancers e.g. ^4-6 7 8^. A number of signalling pathways have been linked with some of these molecules including activation of FAK, TGFβ-SMAD2/3 signalling, PDGF/PDGFR signalling, Wnt and Hh/GLI signalling ^4,9-12^, however it is uncertain if there is a common regulatory mechanism that links them.

One of the key stromal cell types, the activated fibroblast, is a major producer of tumor ECM ^13-16^. These resident or infiltrating cells acquire a phenotype most often characterised by expression of vimentin, alpha smooth muscle actin (αSMA), fibroblast activation protein (FAP), fibroblast-specific protein (FSP), or yes-associated protein (YAP) ^17-19^. Previously we reported a strong positive correlation between the density of αSMA+ and FAP+ stromal cells, and the degree of disease in metastatic HGSOC ^9^. Signalling pathways associated with the activation of αSMA+ fibroblasts have included, most notably, TGFβ as well as and Hh ^18,16,20,21^.

The main goal of this study was to identify cells and signalling pathways regulating production of the six upregulated MI molecules and then build a human multi-cellular model replicating this disease process. We first characterised biopsies from human HGSOC omental metastases and used *in silico* analysis of HGSOC samples to generate hypotheses. We then isolated early passage and primary cell cultures from biopsies for validation and to build an informed novel 3D tri-culture HGSOC model. We report that TGFβR and Hedgehog (Hh) signalling are important regulators of the six upregulated MI molecules that are mainly produced by αSMA+/FAP+ omental fibroblasts, and that cross-talk between these two pathways supports initiation and maintenance of this activated phenotype. Moreover, our novel human HGSOC model replicated some key features found in HGSOC omental biopsies and allowed us to understand clinically relevant regulation of diseased-associated matrisome molecules.

## Results

### Stromal and malignant cells diversely produce disease-associated matrix molecules

We analysed thirty-six omental biopsies with a spectrum of tissue remodelling and disease involvement, all from patients with Stage 3-4 HGSOC. Tissues were assigned a disease score based on area of tissue remodelled with disease-associated stroma and malignant cells (Fig. 1a) as previously described ^3^. Density of PAX8+ cells, a marker of the malignant cells, correlated strongly (*r* = 0.874) with disease score, as expected (Fig. 1b, SFig. 1a). All omenta studied were from patients with confirmed metastases but malignant cells were only visible in 26/36 biopsies.

**Figure 1:**
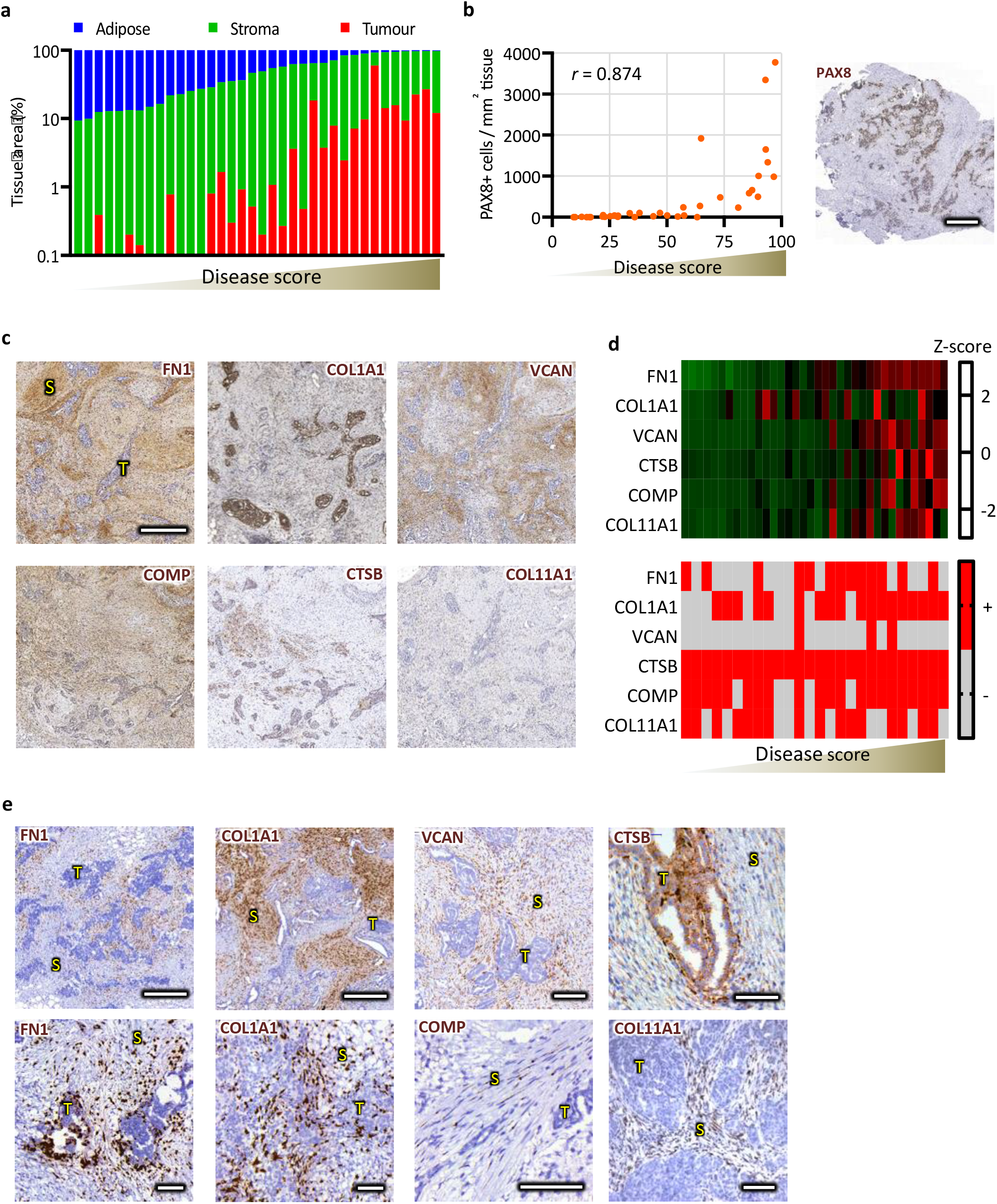
Tumour-matrix proteins are diversely produced by stromal and malignant cells. (**a**) Disease score vs. tissue area (log scale) for 36 stage 3-4 HGSOC patient omental samples correlated with (**b**) PAX8+ malignant cells stained via IHC. (**c**) IHC for FN1, COL1A1, VCAN, COMP, CTSB, COL11A1. All showed positive correlations with disease score and heterogeneous malignant cell staining. (**d**) Z-score heat map (top) of matrix area for all samples and a binary heat map (bottom) displays tumour positivity, red, or negativity, grey, for matrix molecules. (**e**) RNAscope *in situ* hybridisation for the six matrix molecules (brown). Spearman’s rank correlation coefficient, *r*. In panels (**c**) and (**e**), S is stroma and T is tumour. Scale bars are (**b**) 1000μm, (**c**) 200μm and (**e**) (top row COL1A1 and FN1) 500μm, VCAN 200μm, the rest are 100μm.

Matrisome protein density was measured by IHC in tissue sections and quantified with Definiens Tissue Suite® software. Density of all six upregulated-MI molecules increased significantly with disease progression (Fig. 1c-d, SFig. 1b). FN1 had the strongest correlation (*r* = 0.969) with disease score, the greatest average density and was primarily located throughout the stroma, but was also found in malignant cells in 50% (13/26) of biopsies. VCAN was largely confined to stroma with low malignant cell positivity (3/26 biopsies). COMP was present in both stroma and most malignant cells (21/26). CTSB was present in malignant cells of all samples with additional positivity in dense aligned stromal borders. COL11A1 was most common in stroma adjacent to malignant cells or with high cell alignment and there was malignant cell positivity in 16/26 biopsies. COL1A1 had the weakest correlation with disease score and was found heterogeneously throughout the stroma and malignant cells (18/26 biopsies). Figure 1d summarises the pattern of malignant cell positivity for the upregulated-MI molecules.

To further study origin of the upregulated-MI molecules, we conducted RNAscope *in situ* hybridisation on highly-diseased tissue sections (Fig. 1e). Consistent with IHC data, *FN1* and *COL1A1* were expressed in stroma adjacent to malignant cells and in some malignant cells. *VCAN* and *COMP* were expressed mostly in stromal cells with elongated morphology and bordering malignant cells. A minority of malignant cells had *VCAN* expression, however, in contrast to IHC, no *COMP* expression was observed in malignant cells. *CTSB* was expressed in all malignant cells and in stroma at some malignant cell:stromal borders. *COL11A1* was expressed in stroma adjacent to malignant cells or where stromal cells appeared elongated and there was expression in some malignant cells.

In summary, these data support and extend our previous RNAseq and proteomic results in HGSOC confirming positive correlations of the upregulated-MI molecules with disease score and identifying spatial location and cellular origin.

### The prognostic matrisome molecules associate with TGFβ and Hh signalling

Our next aim was to identify common regulatory pathways for the upregulated MI molecules. We first integrated protein-protein interaction and signalling pathway data from public databases using PathwayLinker.org ^22^. Figure 2a illustrates the interaction network of the upregulated MI molecules along with their first neighbour interactors. There was at least one direct or indirect interaction linking *FN1, VCAN, COMP, COL1A1* and *CTSB*. Based on the upregulated-MI molecules and their first neighbour interactors, TGFβ (KEGG) signalling pathway was significantly overrepresented (*p* = 2.2e-09), along with ECM (KEGG), PDGF (Reactome), Actin (KEGG), and P53 (KEGG), (Supplementary Table 1).

**Figure 2:**
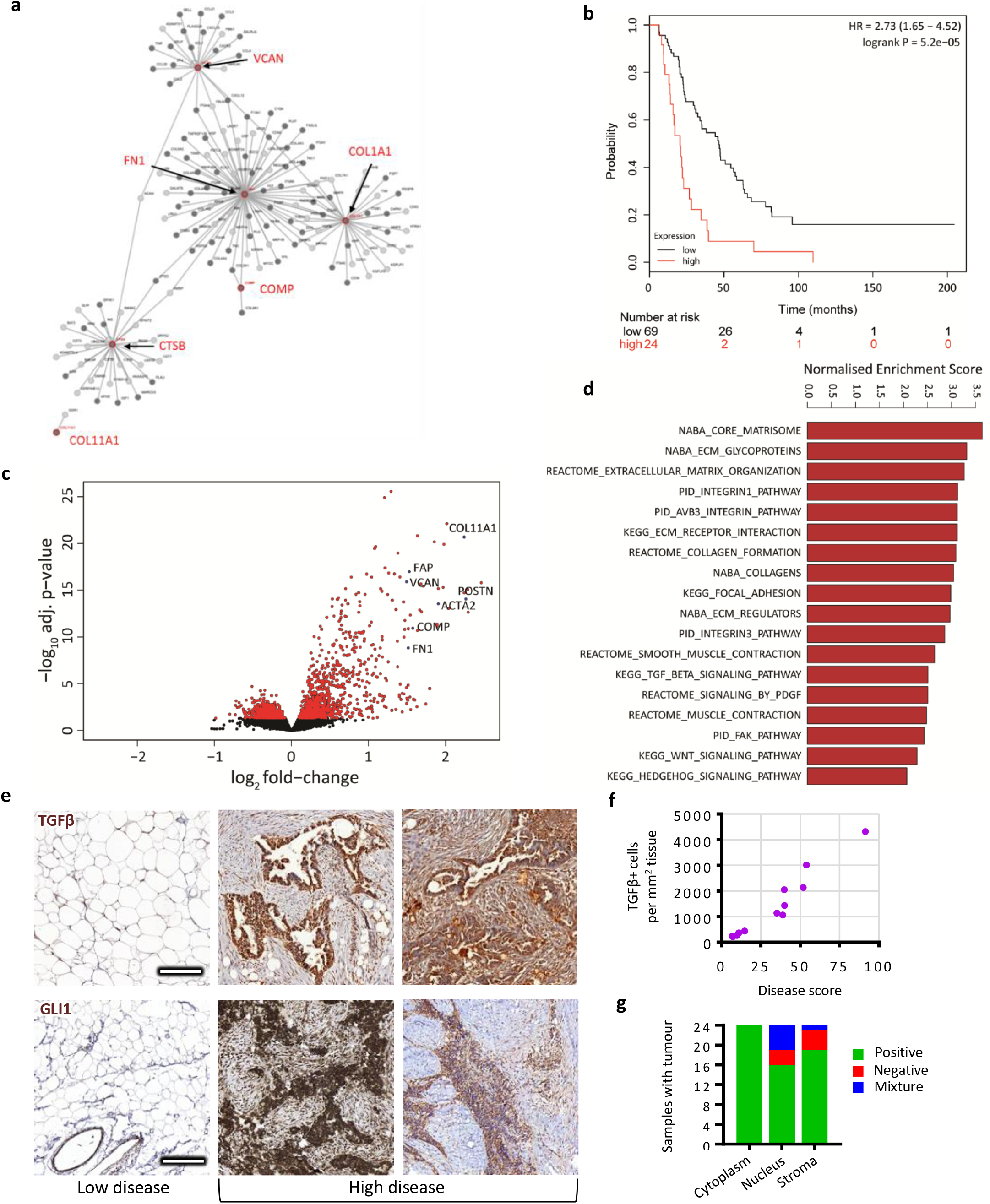
The six upregulated matrix index molecules (FN1, COL1A1, VCAN, COMP, CTSB, COL11A1) associate with TGFβ and Hh signalling. (**a**) Interaction network diagram obtained on PathwayLinker.org for the six upregulated matrix index molecules. (**b**) Kaplan– Meier survival curve with overall survival from the ICGC ovarian dataset divided by high or low average gene expression of the six matrix index molecules. The x-axis is in the unit of months. (**c**) Volcano plot displays comparison of gene expression between high and low groups, highlighting significant upregulation of matrix molecules and stromal activation markers in high group. Red dots indicate adj.P.Value < 0.05. (d) Plot of normalised enrichment score (NES) obtained from GSEA highlights over-expressed pathways in high group compared with low group, including TGFβ and Hh signaling (false discovery rate, FDR < 0.05). (**e**) IHC for TGFβ and GLI1 was performed on biopsies. (**f**) TGFβ IHC was quantified as number of positive cells in stained tissue. (**g**) Samples with malignant cells were identified for populations with GLI1 positivity,negativity or a mixture of both inmalignant cell cytoplasm and nuclei,as well asstromal cells.Scale bars are 200μm.

We next interrogated the ICGC transcriptional dataset of HGSOC biopsies ^23^ looking at association between prognosis and mean expression levels of upregulated MI genes. We found that high average expression of upregulated MI genes associated with significantly worse survival in HGSOC (logrank P = 5.2e-05) (Fig. 2b). Differentially expressed genes in high expression group included periostin, several collagens, osteonectin, and activation markers *ACTA2* and *FAP* (Fig 2c). Gene set enrichment analysis (GSEA) highlighted over-expression of matrisome, ECM, collagen, focal adhesion and smooth muscle contraction pathways (Fig. 2d). Of particular interest was significant enrichment for TGFβ (KEGG), WNT (KEGG), PDGF (Reactome) and Hh (KEGG) signalling (Fig. 2d, Supplementary Table 2). These results suggested that at least five of the upregulated-MI molecules might be co-regulated in HGSOC and that TGFβ signalling is involved.

To investigate this further, we stained tissue sections for TGFβ (Fig. 2e) and found a strong correlation between positive cell density and disease score (Fig. 2f), observing the strongest staining in malignant cells. We also stained tissues for GLI1 (Fig. 2e), a main downstream target of Hh signalling, which has been previously associated with collagen producing αSMA+ phenotypes ^16,20^ and also implicated in TGFβ cross-talk promoting fibrosis ^21,24^. Tissues with high disease score had significant GLI1 positivity in contrast to low disease score tissues that had relatively little. Malignant cells in all biopsies displayed cytoplasmic GLI1, but nuclear GLI1 varied; 16/26 biopsies had total nuclear positivity, 4/26 were totally nuclear negative, and 6/26 were mixed (Fig. 2g). GLI1+ stromal cells were identified in 22/26 of malignant cell biopsies displaying a mixture of cytoplasmic and nuclear positivity.

### Malignant cells upregulate TGFβ3 and heterogeneously express matrisome molecules

Our next aim was to build 2D and 3D *in vitro* human cell models to allow us to validate and extend our findings. First, we investigated two HGSOC malignant cell lines for suitability in such models. G164 and AOCS1 were established in our lab from patients with omental metastases and kept at low passage number. The original tissue biopsies for both cell lines showed malignant cell PAX8 positivity and cell lines cultured *in vitro* maintained PAX8 nuclear positivity (Fig. 3a). We characterised these HGSOC cells by RNA-sequencing. Unsupervised clustering using principle component analysis (PCA) illustrated significant transcriptional differences between the two cell lines with the first principle component accounting for more than 89% of the difference (Fig. 3b). However, within the same cell line there was relatively little variation between monolayer and spheroid culture. GSEA highlighted significant canonical pathway differences between the two cell lines; notably, AOCS1 had enriched Hh-GLI signalling and G164 was enriched in transcriptional activity of SMAD2/3/4 and signalling by TGFβR complex (Fig. 3c). The differences in Hh signalling between AOCS1 and G164 were confirmed by analysing GLI1 expression using qRT-PCR (Fig. 3d) and IF (Fig. 3e). In addition, cell proliferation, which Hh is known to affect, was significantly reduced in AOCS1 with a GLI1/2 inhibitor, GANT61, ^25^ (SFig. 2a-c) but not in G164 (SFig. 2d). Use of a TGFβR inhibitor, SB431542, ^26^ on the G164 cell line attenuated cell contraction of gels and also cell migration (SFig.2e,f), both processes associated with activated TGFβR signalling, but did not affect viability (SFig.2g). RNA sequencing and qRT-PCR detected five of the six upregulated-MI molecules (mRNA for COMP was not present) in the cell lines but expression was heterogeneous (Fig. 3f and SFig. 2h). We stained biopsy sections for differentially expressed matrix molecules identifying malignant cell positivity for COL11A1 in AOCS1 and FN1 in G164 (Fig. 3g). IF on *in vitro* monocultures displayed strong intracellular COL11A1 staining organised into fibrils in AOCS1 and significant deposition of FN1 by G164s (SFig. 2i) replicating tissue staining.

**Figure 3:**
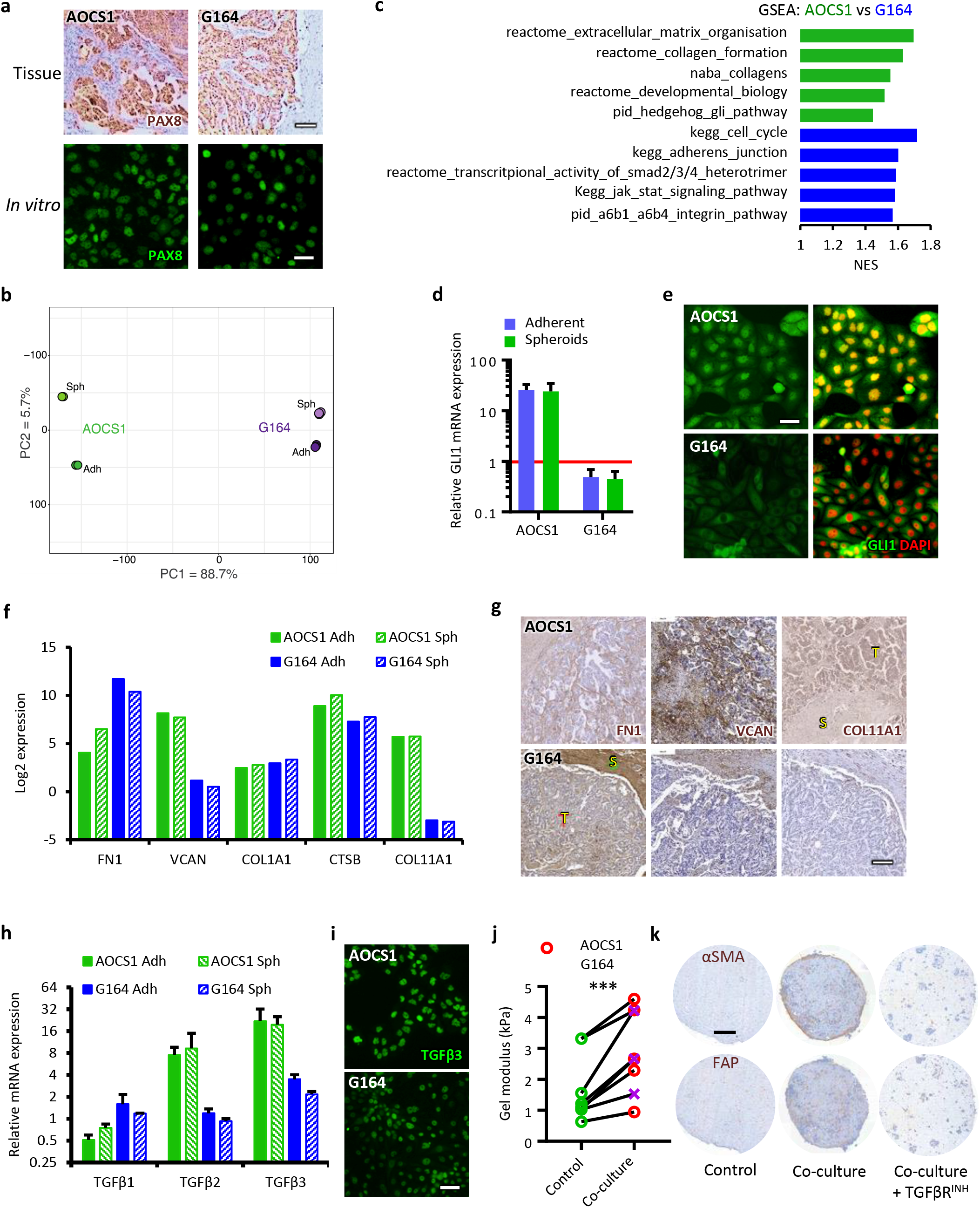
HGSOC malignant cells have heterogeneous signaling but all secrete TGFβ. (**a**) To identify malignant cells, HGSOC tissue sections and cells expanded *in vitro* were stained for PAX8 via IHC or IF respectively. (**b**) RNA sequencing was performed on AOCS1 and G164 malignant cell lines (N = 2) for adherent (Adh) and spheroid (Sph) cultures and transcriptomic expression was analysed using PCA. (**c**) GSEA was performed on transcriptomic data and normalised enrichment scores for AOCS1 (green) vs G164 (blue) are illustrated (p < 0.1). (**d**) qRTPCR was performed for GLI1 on AOCS1 and G164, barplot illustrates relative expression levels of GLI1 normalised to expression in normal fallopian tube cells, FT318-WT (red line) (e) GLI1 IF (f) RNASeq log2RPKM gene expression of matrix molecules in AOCS1 and G164 cultures (COMP not detected) and (g) IHC of human biopsy sections from patients AOCS1 and G164 for FN1, VCAN, and COL11A1. (**h**) TGFβ expression via qRTPCR for malignant cells *in vitro* (N = 3), normalised to FT318-WT expression and (**i**) IF images of TGFβ3. (**j-k**) OFs were cultured in 3D COL1 gels alone (control) or with (co-culture) AOCS1 or G164 cells for 7-14 days with or without TGFβR^INH^, then (**j**) compressed to determine gel modulus (D7); each point represents mean of 2-4 gels (n=2-4) per OF donor (N = 8), p < 0.001 (two-way, paired t test) and (**k**) IHC (D14) performed on fixed sections of AOCS1 co-cultures for αSMA and FAP.Scalebars(**a** (bottom),**e**,**i**) are 50μm,(**a**(top)**g**,**k**) are 250μm

Expression of the three TGFβ-isoforms was analysed in malignant cells and compared to a non-malignant cell control, wild-type immortalised FT318 fallopian tube surface epithelial cells (Fig. 3h). Malignant cells expressed all three TGFβ isoforms with little difference between adherent and spheroid cultures, but TGFβ3 was the most highly expressed relative to FT318. TGFβ3 protein was also confirmed by IF in both cell lines (Fig. 3i). To test the influence of malignant cell-secreted TGFβ on fibroblast activation, we co-cultured malignant cells with primary omental fibroblasts (OFs) in collagen gels. All co-culture gels had a significantly greater gel modulus (stiffness) compared with respective fibroblast controls (Fig. 3j), while malignant cells grown alone did not alter gel modulus (SFig. 2j). In co-cultures, αSMA, FAP and eosin staining were all increased in OFs, but expression was attenuated using the TGFBR inhibitor (Fig. 3k, SFig. 2k). Interestingly, we were unable to detect expression of any Hh ligands in either cell line.

These results highlight heterogeneity in cell-signalling and matrisome expression in HGSOC malignant cells but they also reveal a commonality in upregulation of TGFβ, which activates OFs via TGFβR signalling.

### TGFβ3 stimulates activation, GLI1 expression and diseased-matrix production in omental fibroblasts

Having identified TGFβ as a stimulator of fibroblast activation, we next looked at associations between fibroblasts and malignant cells in HGSOC omental biopsies. Density of αSMA+ or FAP+ cells correlated with density of PAX8+ malignant cells (*r* = 0.833 and *r* = 0.814 respectively) (Fig. 4a) and staining was highest in stroma adjacent to malignant cells (SFig. 3a). We pseudo coloured and overlaid consecutive αSMA and FAP tissue images (Fig. 4b) identifying an αSMA+/FAP+ stromal phenotype located primarily at malignant cell borders where the densest matrix and GLI1 staining were previously seen.

**Figure 4:**
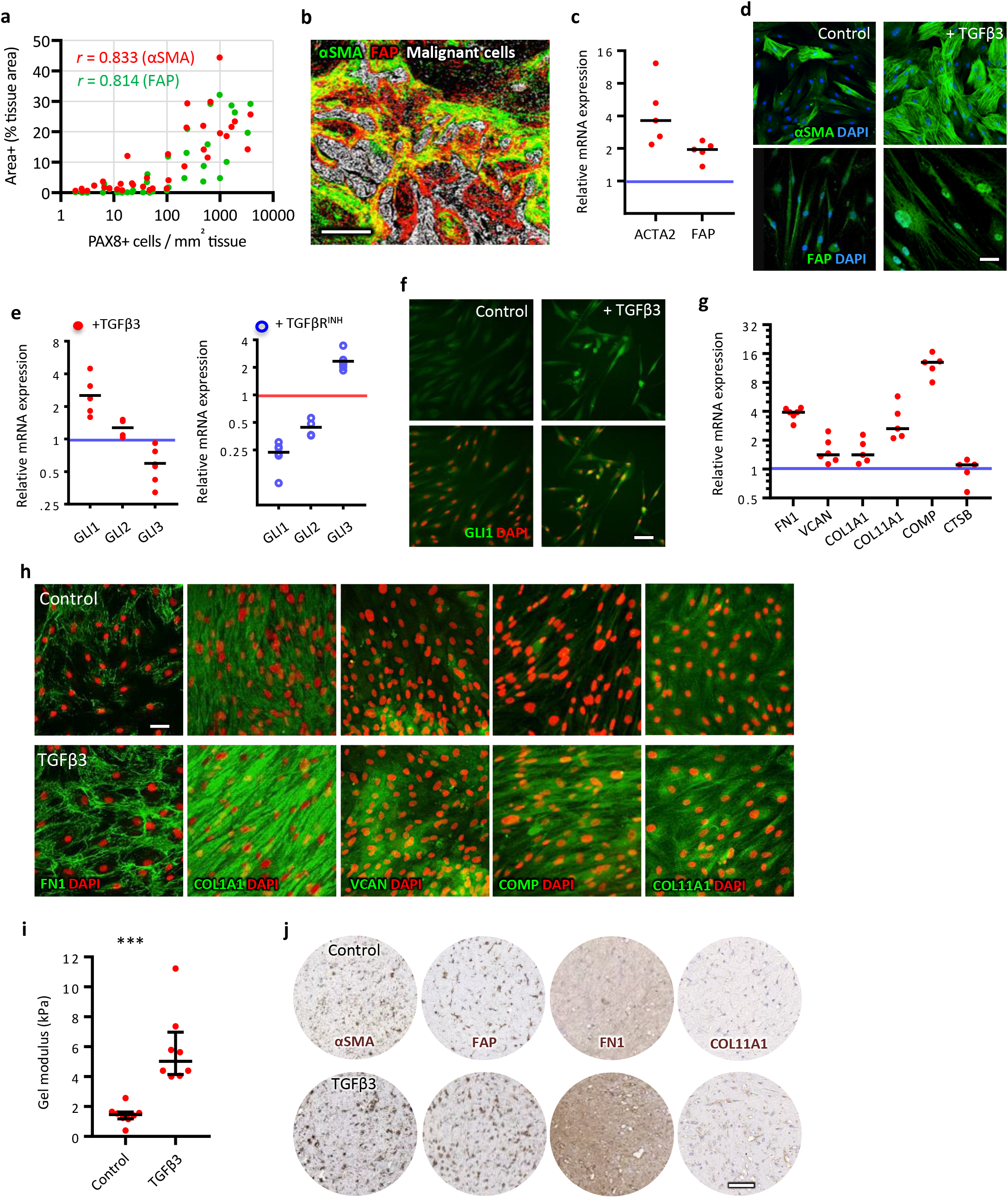
TGFβ3 stimulates αSMA, FAP and GLI1 expression and tumour-matrix production in omental fibroblasts. (**a**) IHC identified FAP or αSMA positive cells; both correlated positively with PAX8+ cell density. (**b**) IHC images of FAP and αSMA were pseudo colored and overlaid using ImageJ to highlight double positive cells (yellow). (**c**) TGFβ3 treatment of L-OFs caused upregulation in ACTA2 and FAP mRNA, measured by qRTPCR, across five donors (n=3, N=5) and (**d**) increased αSMA and FAP IF. (**e**) GLI1 mRNA expression was upregulated in TGFβ3-treated L-OFs (5 donors) and downregulated with SB431542 (TGFβR^INH^) (4 donors). (**f**) IF GLI1 staining shows representative images of TGFβ3-treated and untreated L-OFs. (**g**) Five of the six matrix molecules were upregulated in TGFβ3-treated L-OFs at mRNA level. (**h**) Representative IF staining for the five upregulated molecules. (**i**-**j**) L-OFs were cultured in COL1 gels for seven days; TGFβ3-treatment increased (**i**) gel modulus (n=2-3, N=3, two-way t-test, *p* < 0.001) and (**j**) expression of αSMA, FAP, FN1 and COL11A1 visualised via IHC. Scale bars are (**b**) 200μm and (**d, f, h**) 50μm and (**i**) 500μm. Blue and red lines represent control (no TGFβ3 or no TGFβR^INH^ respectively) expression.

When we isolated OFs from HGSOC omental biopsies we observed a range of activation states in culture, which we categorised before use in experiments as either low (L-OFs) or high (H-OFs), defined by cell morphology and level of expression of F-actin and αSMA stress fibres (SFig. 3b). Regardless of initial activation state, treating OFs with TGFβ3 increased *ACTA2* and *FAP* mRNA expression on average 4-fold and 2-fold respectively (Fig. 4c), increased IF staining for both proteins (Fig. 4d, SFig. 3c) as well as the proportion of αSMA+/FAP+ cells (SFig. 3d). TGFβ3-treatment also promoted F-actin fibre formation, FSP and YAP nuclear expression (SFig 3e) in L-OFs, although differences in the latter two appeared small. In Figure 2e we identified GLI1 positive stromal cells in HGSOC tissue. Hh activation can promote collagen-producing myofibroblasts in some fibrotic diseases ^27^ and recently αSMA+/GLI1+ mesenchymal cells have been identified ^28^. TGFβ3-treated OFs had on average a 2.5-fold increase in GLI1 expression and a 2-fold decrease in GLI3 expression, typically considered a Hh pathway repressor ^29^, while SB431542 reversed this trend (Fig. 4e). GLI1 IF showed weak cytoplasmic staining in L-OFs while TGFβ3-treated L-OFs had stronger cytoplasmic and positive nuclear staining (Fig. 4f).

Next we analysed expression of upregulated-MI molecules in L-OFs and found that TGFβ3 upregulated mRNA expression of *FN1, VCAN, COL1A1, COL11A1*, and *COMP* (Fig. 4g) and promoted matrix deposition and organisation (Fig. 4h). Initially, there was little VCAN or COMP in confluent untreated L-OFs but TGFβ3 induced widespread deposition of both proteins. Treated-cells contained denser FN1, COL1A1 and COL11A1 fibres with greater alignment. Of particular note, COL11A1 was organised into intracellular fibres similar in appearance to microtubules.

Tissue stiffening and contraction occurs during remodelling in fibrosis-associated diseases and is caused largely by myofibroblasts ^26,30^. When we cultured L-OFs in 3D collagen gels for 7 days, TGFβ3-treatment caused ∼ 4-fold increase in gel modulus compared with untreated gels (Fig. 4i). Gels seeded with H-OFs formed stiffer gels than L-OFs but modulus was significantly reduced when treated with SB431542 (SFig. 3f). Cells in TGFβ3-treated gels had denser αSMA, FAP, FN1 and COL11A1 staining compared with controls (Fig. 4j).

Collectively these experiments showed that TGFβ3 promotes an αSMA+/FAP+ contractile OF phenotype associated with an upregulation of GLI1 and increased deposition of five of the six upregulated-MI molecules.

### TGFβR and GLI1/2 inhibitors downregulate αSMA, GLI1 and matrix molecules in OFs

We next asked if GLI1 played a downstream role in TGFβR pathway activation in H-OFs (Fig. 5) and TGFb3-activated L-OFs (SFig. 4). Inhibitors of TGFBRI, SB431542 ^25^ or Gli1/2, GANT61 ^26^, both reduced αSMA stress fibres in H-OFs and induced a morphology shift from a relatively large spread cell to a smaller fusiform cell (Fig. 5a). Both inhibitors reduced *ACTA2* expression between 4-5 fold, but there was less effect on FAP (Fig. 5b, SFig. 4a). Combination treatment of the inhibitors caused a synergistic effect on *ACTA2* resulting in >10-fold down-regulation. Both inhibitors decreased the proportion of αSMA+/FAP+ cells with the greatest effect induced by the inhibitor combination (Fig. 5c, SFig. 4b). IF confirmed αSMA stress fibres decreased with inhibitor combination, while again there was less effect on FAP (Fig. 5d). Both inhibitors down-regulated GLI1 3-5 fold and upregulated GLI3 2-3 fold (Fig. 5e, SFig. 4c) while IF showed reduced cytoplasmic and nuclear GLI1 in H-OFs treated with the inhibitor combination (Fig. 5f).

**Figure 5:**
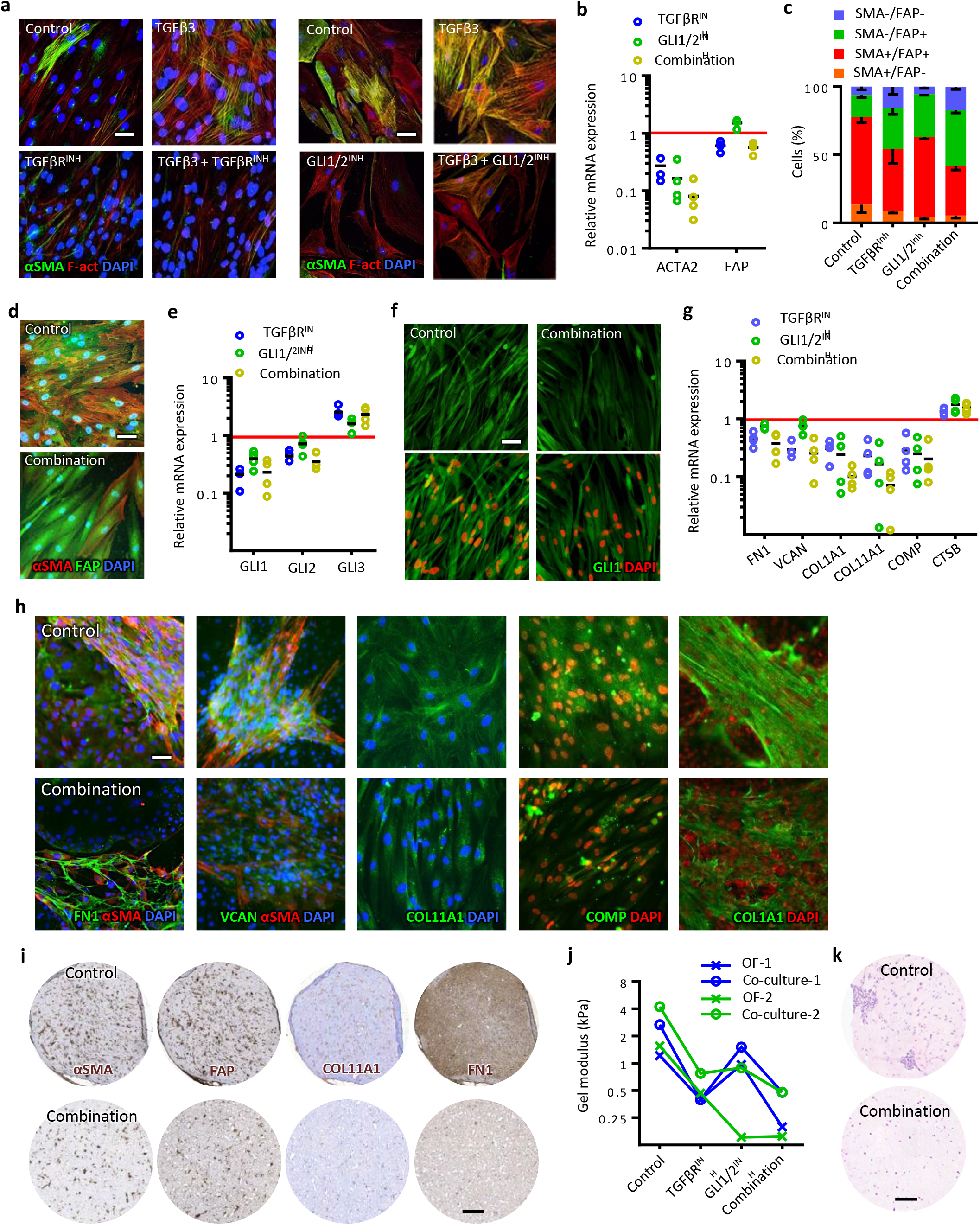
TGFβR and GLI1/2 inhibitors downregulate αSMA, GLI1 and tumour-matrix in OFs. (**a**) IF for αSMA and F-actin in L-OFs treated with TGFβ3 ± TGFβR^Inh^ (SB431542) or ± GLI1/2 (GANT61) inhibitors. (**b-k**) OFs with high activation (H-OFs) were treated with TGFβR^Inh^ or GLI1/2^Inh^, or combination; (**b**) gene expression was measured by qRTPCR for ACTA2 and FAP (N = 4 donors). (**c**) Cells were analysed via flow cytometry for % cells αSMA+ and/or FAP+. (**d**) Activated OFs were visualised via IF for αSMA and FAP ± Inhibitor combination. (**e**) GLI1/2/3 mRNA expression analysed via qRTPCR for OFs and (**f**) GLI1 visualised via IF in activated OFs or with combination. (**g**) mRNA expression of matrix molecules were analysed via qRTPCR for treated or untreated OFs (n = 4 donors). (**h**) IF visualised FN1, VCAN, COL11A1, COMP and COL1A1, and αSMA in activated OFs with or without combination. (**i**) IHC of control and combination-treated OF-gels for αSMA, FAP, COL11A1 and FN1. (**j-k**) OF-only gels or AOCS1 co-cultures were grown for 7 days with or without inhibitors and (**j**) gel modulus was measured (mean of 3 gels per donor, N=2) and (**k**) fixed sections stained for H&E. Scale bars (**a, d, f, h**) are 50μm, (**I, k**) are 500. Red lines represent control (no TGFβR^INH^) expression.

SB431542 reduced *FN1, VCAN, COL1A1, COMP* and *COLL11A1* mRNA while GANT61 reduced *COL1A1, COMP* and *COL11A1* (Fig. 5g, SFig. 4d), implying that FN1 and VCAN are not regulated by Hh. Overall the inhibitor combination was most effective at downregulating collagen matrix mRNA than each individual inhibitor. Figure 5h shows representative IF in 2D H-OF cultures for the five molecules down regulated at mRNA level molecules confirming that the effect of the inhibitor combination is replicated at protein level. There was almost complete absence of VCAN and COMP; fibres of FN1 and COL1A1 were less and disrupted; and COL11A1 had lost fibrous structure. When H-OFs were grown in 3D collagen gels the inhibitor combination reduced density of αSMA, FAP, FN1 and COL11A1 (Fig. 5i). In contrast to OFs, there was relatively little inhibitor effect on matrix expression or organisation in AOCS1 malignant cells (SFig 5a-b). However, in G164 malignant cells, there was a significant effect of the TGFβR inhibitor on FN1 mRNA and protein (SFig. 5c-d), following the trend seen in OFs, and this translated to 3D cultures whereby spheroid growth was also reduced (SFig. 5e). In addition, we observed a marked reduction in *TGF*β*2* and -β*3* expression in AOCS1 with Hh inhibitor GANT61 and a 10-fold reduction in *TGF*β*2* expression in G164 with TGFBR inhibitor SB431542 (SFig. 5f-g).

We next co-cultured malignant cells with OFs in 3D gels and measured inhibitor effect on gel modulus. Co-cultures treated with either inhibitor alone had decreased modulus and overall the combination was the most effective (Fig. 5j), also reducing the size of cell clusters in the gels (Fig. 5k). Viability of co-cultures was unaffected by inhibitors (SFig. 5h).

In summary, we found that TGFβ stimulates GLI1 promoting an αSMA COL1A1/COL11A1/COMP rich matrix, however, Hh signalling appeared to have little involvement with VCAN or FN1 production. We also observed a synergy with TGFβR and GLI1 inhibition resulting in the inhibitor combination being most effective in attenuating activation, contraction and matrix production. There were also regulatory differences in the two pathways between malignant cells and OFs.

### A novel 3D tri-culture replicates some key features found in HGSOC biopsies

In order to conduct pre-clinical studies of therapies that could target the poor prognostic MI molecules we require human 3D models that recreate more of the complexity and interactions in the tumor microenvironment. Adipocytes are the major cell type in normal omentum and act as a source of energy for developing metastases ^31^. We hypothesised that adipocytes could provide a relevant biological substrate for longer-term malignant cell and fibroblast co-culture allowing for the study of matrisome molecule regulation and also that such tri-cultures would also more closely resemble the omental tumor microenvironment. We isolated mature adipocytes from omental digests (Fig 6a) and confirmed there were viable unilocular cells. We seeded adipocytes into low-weight (0.1w%) COL1 gels in 96 well plates, enabling adipocytes to rise up forming a compact mm-sized layer (Fig 6b). Figure 6c shows H&E images of an adipocyte gel after 14 days of culture compared to normal human omentum. We assessed viability and perilipin-1, a marker of mature adipocytes, over 21 days. Although there appeared to be some decline in viability/live stain there was very little dead staining at day 21 (Fig 6d) and perilipin-A levels remained relatively constant (Fig 6e). IHC for MI molecules revealed low levels of MI molecules in the adipocyte cultures (SFig. 6a).

**Figure 6.**
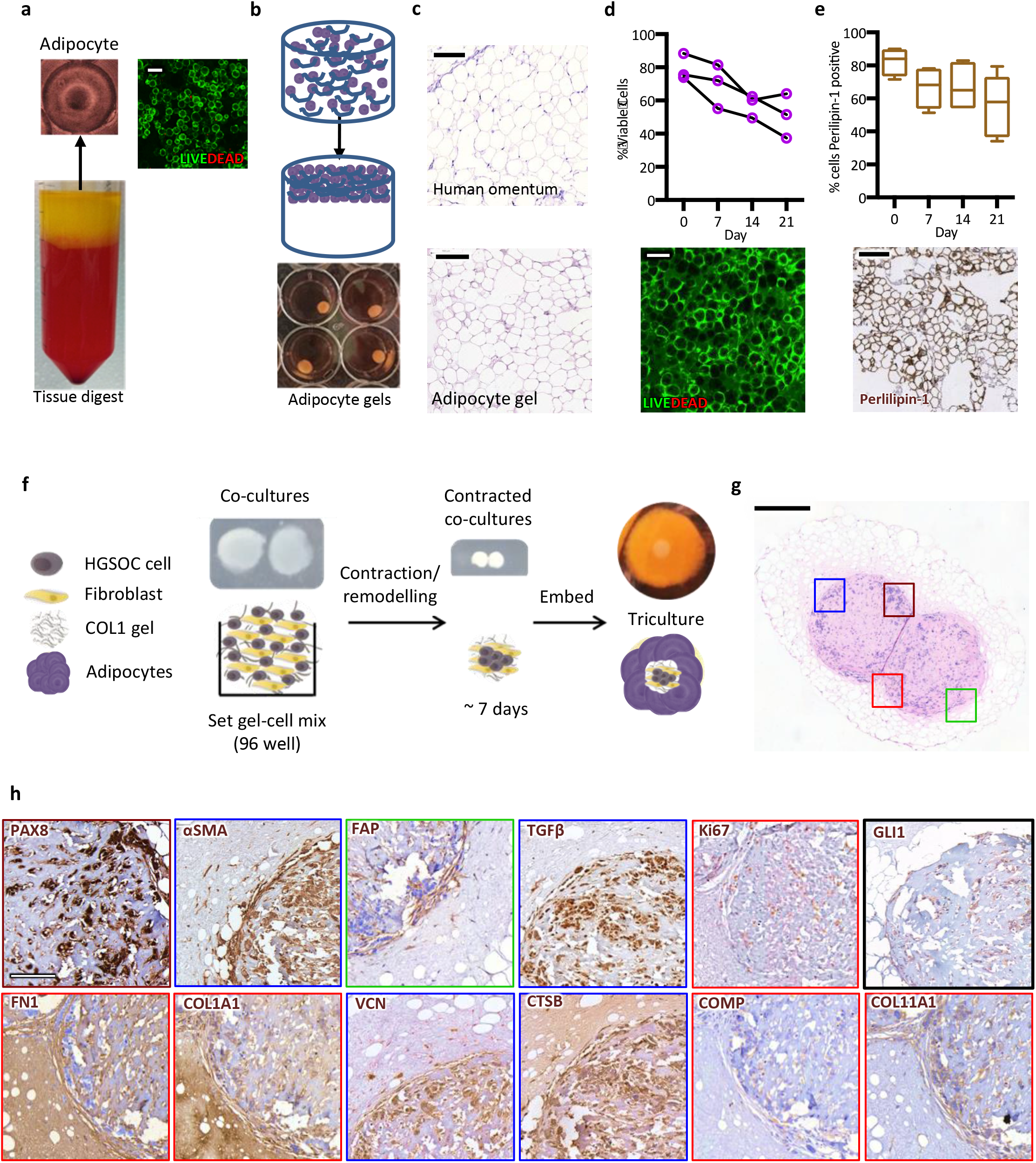
Omental adipocytes provide a physiological substrate for a HGSOC tri-culture model. (**a**) Fatty layer on top of omental digest supernatant contains viable adipocytes, assessed by IF LIVE/DEAD assay. (**b**) Adipocytes are mixed with collagen gel solution (0.1w%), seeded into 96-well plates, left 5min at RT allowing cells to float upwards, before incubating at 37°C for 45min, after which gels can be carefully handled (gels transferred to 24-well plates). (**c**) H&E gel sections have similar appearance to normal omentum. (**d-e**) Adipocyte gels tested for viability via IF LIVE/DEAD assay and sectioned and stained via IHC for Perilipin-1 (days 0, 7, 14 and 21); data are (**d**) mean of 3-5 images per donor (N = 3) and (**e**) median with interquartile range (N = 5). (**f**) Schematic shows constituents and cells used to assemble and grow tri-cultures. (**g-h**) AOCS1 tri-cultures grown for 21 days, then fixed, sectioned and stained for H&E and via IHC, coloured squares in (**g**) mark location of coloured borders for IHC (**h**). Scale bars (**a, c, d**) are 100μm, (**e, h**) are 200μm, (**g**) is 500μm.

To establish tri-cultures, we inserted seven-day co-cultures (providing time for gel remodelling) directly into the middle of adipocyte gels and then incubated for a further 14 days in free-swelling conditions (Fig. 6f). This time period was sufficient for remodelling of the adipose tissue, and generation of cell markers and all upregulated MI molecules (Fig. 6g-h, SFig. 6b). PAX8 staining identified malignant cells, while FAP and αSMA staining identified activated fibroblasts located mainly at borders between malignant cells and remodelled adipose tissue (Fig. 6h) as we had observed in biopsies. Both malignant cells and stromal cells were proliferating at 21 days demonstrated by Ki67 staining (Fig. 6h). Malignant cells were positive for TGFβ and GLI1 positivity was present in malignant and stromal cells. All six upregulated-MI molecules were found in the stroma and some in malignant cells (Fig. 6h). We conclude that tri-cultures support malignant cell and fibroblast growth and replicate some of the key features that we have found in patient biopsies.

To further confirm that the tri-cultures were a valid model to study regulation of MI molecules and that the addition of adipocytes better replicated the human omental TME, we conducted RNAseq analysis of the adipocyte gels, G164-OF collagen gel co-cultures and the G164 tri-cultures (Fig 7 a-d). The tri-cultures had the most complex matrisome gene transcription signature (Fig. 7a) and showed a significant enhancement of ECM, adhesion, collagen fibril organisation and also cell migration signatures compared to the G164-OF co-cultures (Fig 7b, Supplementary Table 3). Twenty-one of the 22 MI molecules originally identified by us in HGSOC omental metastases ^3^ could be sufficiently detected by RNAseq allowing us to calculate the MI of adipocyte-only gels, G164-OF cultures and tri-cultures. Tri-cultures had an increased MI compared with adipocyte-only gels and co-cultures (Fig. 7c) with values similar to diseased omental metastases from HGSOC patients ^3^. The cluster dendrogram also shows a clear separation between adipocyte cultures, co-cultures and tri-cultures (Fig. 7d).

**Figure 7.**
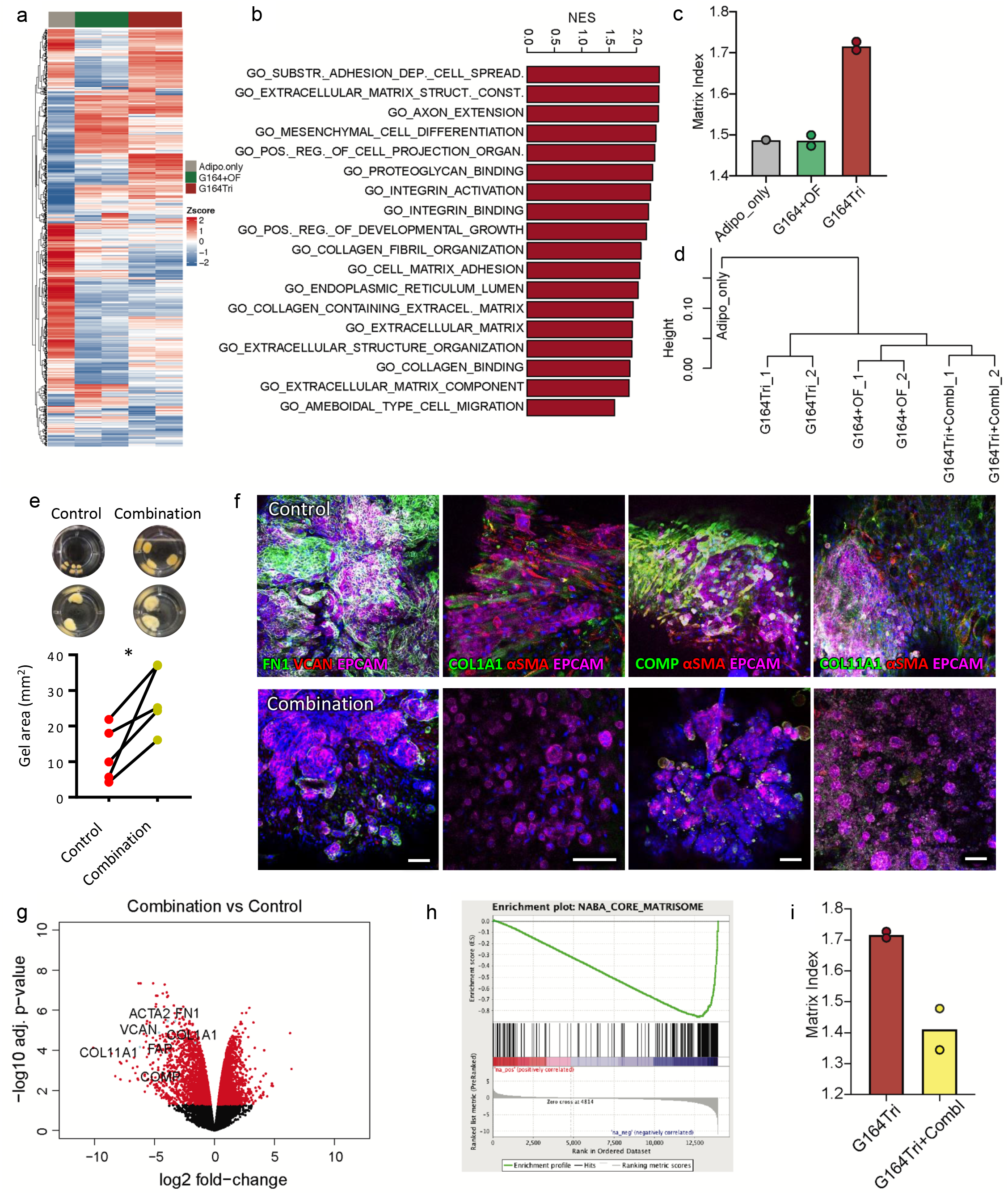
TGFβR and GLI1/2 inhibitor combination reduces tumour matrix expression in HGSOC tri-cultures. RNAseq was performed on adipocyte-only gels (Adipo_only), control tri-cultures (G164Tri), G164+OF co-cultures and G164Tri treated with inhibitors combination (G164Tri+CombI). (**a**) Heatmap of all matrisome genes detected in Adipo_only, G164+OF and G164Tri. (**b**) GSEA was perfomed on differentially expressed genes in G164Tri vs G164+OF, barplot indicates normalized enrichment scores (NES) for the indicated Gene Ontologies (p < 0.05). (**c**) Matrix Index was calculated from the RNAseq data across the samples. (**d**) Hierarchical cluster analysis of transcriptomes for indicated samples. (d) After 21 days of culture, gel area of control and combination-treated tri-cultures were measured from images (inserts show 2 different experiments); data are mean of 2-4 gels per experiment (N = 5), p < 0.05 (two-way, paired t test). (**f**) IF on tri-cultures for EPCAM, αSMA, FN1, VCAN, COL1A1, COMP, COL11A1, and DAPI (blue), scale bars are 100μm. G164Tri were treated with inhibitor combination and subjected to RNAseq (**g**) Volcano plot used to visualise changes in expression between G164Tri treated with inhibitors (G164Tri+CombI) and control G164Tri. Red dots indicate adj.P.Value < 0.05. (**h**) GSEA performed on the differentially expressed genes between G164Tri+CombI and G164Tri; Enrichment plot derived from GSEA of RNA sequencing data of G164 tri-cultures. (**i**) Matrix Index for G164Tri and G164Tri+CombI. For RNA sequencing, 2 gels were pooled per sample, Adipo_only (n = 1), G164Tri and G164Tri+CombI (n = 2).Scale bars are 100μm.

### TGFβRI and GLI1/2 inhibitor combination reduces matrisome production in tri-cultures

Having shown that MI levels were comparable to the biopsies, and implicating TGFβR and Hh signalling in regulation of the upregulated-MI molecules, we tested their inhibitors in the tri-cultures. We used the inhibitor combination as we saw previously that it was most effective at reducing OF activation and matrisome molecules. The inhibitor combination did not affect total cell viability (SFig. 7a) but significantly reduced adipocyte gel contraction in all tri-cultures (Fig. 7e and SFig. 7b) and reduced remodelling or malignant cell invasion of adipocytes (SFig. 7c). Confocal microscopy showed a noticeable reduction in all six upregulated-MI molecules imaged in tri-cultures treated with the inhibitor combination (Fig. 7f and SFig 7d). In controls, G164 cells formed large colonies surrounded by activated fibroblasts but with the inhibitor combination, malignant colonies were significantly smaller and widely dispersed. In treated AOCS1 tri-cultures, matrix molecules and activation markers were also reduced compared with controls (Fig. 7f and SFig 7d).

We also conducted RNAseq on G164 tri-cultures with the inhibitor combination. Unsupervised clustering of the data showed that the inhibitor-treated cultures segregated separately from the control tri-cultures (SFig. 7e). Expression levels of fibroblast activation markers and the six upregulated-MI molecules, including *CTSB*, were significantly reduced by the inhibitor combination (Fig. 7g). GSEA showed significant downregulation of pathways associated with matrisome, ECM, collagens, as well as TGFβ and Hh signalling (SFig. 7f,g, Fig. 7h, Supplementary Table 4). The overall MI was reduced to the level of the adipocyte gels or co-cultures (Fig. 7i). These experiments demonstrate that this novel tri-culture model replicates key features of the omental HGSOC tumor microenvironment, especially matrisome components from the Matrix Index signature. Moreover, we can use this model to investigate regulation of tumor-associated components.

## Discussion

We recently published a multi-level analysis of developing HGSOC metastases ^3^. One of the significant findings was a pattern of 22 matrisome genes we termed the Matrix Index (MI) that significantly changed with disease progression and was highly prognostic in ovarian and twelve other solid human cancers. Six of these molecules were significantly upregulated with disease progression and sixteen downregulated. In the present work, we have shown that expression levels of the six upregulated-MI molecules themselves predict poor prognosis in HGSOC and associate with an activated αSMA+/FAP+ fibroblast phenotype regulated by TGFβR activity and Hh signalling.

We used the knowledge gained from our previous analysis of HGSOC metastases, further studies on HGSOC biopsies, *in silico* analysis, and *in vitro* cultures to inform and build a relevant 3D multi-cellular model of the tumor microenvironment. We facilitated sustained production of key matrisome proteins in tri-cultures by cell types also found in patient biopsies and RNAseq analysis demonstrated that several important features of the diseased biopsies, especially related to ECM regulation, cell adhesion and migration, as well as the MI gene expression signature, were enhanced in the tri-cultures. Interestingly, compared to responses with monolayers on stiff plastic, FAP expression was significantly down regulated and CTSB expression was also down regulated with the inhibitor combination in tri-cultures. These differences implicate the importance of a physiologically relevant biomechanical environment and our results show that human 3D multi-cellular models can be useful for studying some aspects of cancer biology.

We demonstrated that TGFβ signalling plays a powerful role in induction of an aggressive fibroblast phenotype, which is responsible for deposition of disease-associated matrix that predict poor prognosis. TGFβ ligands play important roles in development, homeostasis and wound healing and all three known isoforms act through the same receptor signalling pathway ^32^. The TGFβ pathway is widely acknowledged as essential for tumor progression and can play pivotal roles as both a promoter and suppressor of cancer cells. Cancer cells can acquire loss-of-function mutations and lose responsiveness to TGFβ thereby bypassing cell cycle arrest ^33^. TGFβ plays an important role in recruitment and activation of cells of the innate immune system but also acts to supress immune cell functions ^34^. Additionally, TGFβ plays an essential role in regulation of the adaptive immune system and its continued presence can supress T-cell functions and promote pro-tumorigenic phenotypes ^35,36^.

More recently, TGFβ has been recognised for its potential regulatory role in the stromal microenvironment, which in turn plays an important role in tumor progression. In our study we identified that malignant cell lines expressed all three TGFβ isoforms and TGFβ-2/3 ligands were upregulated compared to our non-malignant control with the greatest increase for TGFβ3. TGFβ3 has previously been associated with a set of poor outcome genes in serous ovarian cancer ^10^. We showed that TGFβ3 induced αSMA, FAP and GLI1 expression in OFs and promoted production of five upregulated-MI molecules. Of particular interest is a recently published bioinformatics analysis of pan-cancer transcriptional-ECM regulation in cancer ^37^. Chakravarthy *et al* reported that their ECM signature was linked to TGFβ signalling and was a biomarker of failure to respond to immune checkpoint blockade ^37^. Response to an anti-PDL1 agent in patients with metastatic bladder cancer was also associated with increased TGFβ signalling in patient biopsies ^38^ and treatment of tumor bearing mice with inhibitors of TGFβ or its receptor enhanced response to anti-PD1/PDL-1 therapies ^36 38^.

We have highlighted apparent cross-talk between TGFβR and Hh, both pathways promote αSMA in OFs such that TGFβR stimulates GLI1 signalling leading to increased αSMA expression. FN1 and VCAN were upregulated by TGFβ, in agreement with previous studies, but Hh appeared to have little involvement. However, there was a strong indication that Hh enhances COL1A1, COL11A1 and COMP and that the GLI1/2 inhibitor created a synergistic inhibitory effect when used with the TGFβR inhibitor. Interestingly, the GLI1/2 inhibitor did not influence matrix molecule expression in malignant cells, even in AOCS1, which had enriched Hh-GLI pathway activity, implying that the signalling mechanisms either have a functional loss or are different between cell types. However, GLI1/2 inhibition did reduce proliferation in AOCS1. In addition, expression of Hh ligands was not detected in either HGSOC cell line indicating that GLI1 activation in OFs was not due to malignant cell-secreted Hh ligands. These two separate observations of inhibition synergy and GLI1 stimulation by TGFβ signalling suggest that Hh is likely to be activated by more than one pathway and therefore inhibition of TGFβR activity does not silence all Hh activity. This demonstrates potential for use of GLI inhibitors in cancers that have poor prognostic Hh-stromal signatures and malignant cell Hh activity and there may be added benefit from combination treatment with TGFβR inhibitors. Currently, the most clinically advanced group of Hh inhibitors target smoothend (SMO) ^39^, which is upstream of GLI1. However, there have been cases where malignant cells developed clinical resistance to SMO inhibitors via a number of different mechanisms ^39^ and therefore directly targeting GLI may offer more promise in bypassing SMO-resistant cells.

All six upregulated-MI molecules have previously been linked to tumor progression. In particular, COL11A1 expression is consistently linked with poor prognosis in solid metastatic carcinomas ^8,10,12^. While COL11A1 has been implicated as a specific biomarker of activated fibroblasts ^8^, it can also be expressed in epithelial cells with high metastatic potential ^12^. Indeed, in this study we demonstrated that AOCS1 were highly positive for COL11A1, and there was malignant cell nuclear positivity in >50% of our biopsies. Interestingly, in our cells, COL11A1 was intracellular and highly expressed in cells with active Hh signalling. Use of the inhibitor combination on OFs caused a total breakdown of COL11A1 fibrous structure, in line with loss of αSMA stress-fibres, suggesting a role in forming a stable and contractile phenotype. We believe that this warrants further investigation.

In conclusion, the main drive for creating human models of the tumor microenvironment is to study processes governing disease progression in a more physiologically relevant setting and to aid pre-clinical testing. The multi-cellular model we describe here could be useful for screening compounds that could modify the malignant matrisome that associates with poor prognosis in 13 common human cancers. The most promising candidates could then be tested in mouse models that most closely replicate the human TME, either patient-derived xenografts or models such as our recently published new syngeneic mouse HGSOC models that replicate many features of the human omental TME ^40^.

In our model, TGFβR and GLI inhibitors attenuated fibroblast activation and tumour-associated matrix production while preventing malignant cells from forming large spheroid growths. Therefore, inhibitors of these pathways may have clinical potential, alone or in combination. While we do not know if these processes have the potential to increase malignant cell dissemination due to removal of the physical matrix barrier, it is also possible that this may facilitate better access for cancer-treatments or immune cells. We believe that our novel tri-culture model will be a useful first step in pre-clinical evaluation of therapies targeting dysregulated matrix in human solid cancers and their effect on immune cell access to malignant cells. Moreover, we believe that our work demonstrates the usefulness of using a combination of mono-co- and multi-cellular cultures to understand cell:cell interactions in the tumour microenvironment.

## Methods

### Ovarian Cancer Patient Samples and Study Approval

Patient samples were kindly donated by women with HGSOC undergoing surgery at Barts Health NHS Trust. Tissue deemed by a pathologist to be surplus to diagnostic and therapeutic requirement were collected together with associated clinical data under the terms of the Barts Gynae Tissue Bank (HTA license number 12199. REC no: 10/H0304/14). Each patient gave written informed consent and all tissue used for this study was approved by a UK national review board. Studies were conducted in accordance with the Declaration of Helsinki and International Ethical Guidelines for Biomedical Research Involving Human Subjects (CIOMS).

### RNA *in situ* hybridization

Sections (4μm) of formalin-fixed paraffin embedded (FFPE) human omentum samples were deparaffinised, treated with hydrogen peroxide and boiled in the target retrieval reagent. Sections were dried in ethanol and left at room temperature (RT) overnight. Slides were incubated in protease reagent at 40°C in a HyBEZ Hybridization System (Advanced Cell Diagnostics Inc. USA) followed by incubation at 40°C with the gene-specific probe. The AMP 1-6 reagents were all subsequently hybridized as specified in the manufacturer’s instructions. Labelled mRNAs were visualized using DAB reagent and counterstained using 50% Gill’s haematoxylin. Counterstained slides were dehydrated using 70% and 95% ethanol and cleared in xylene before mounting coverslips using DPX. RNAscope® probes: FN1 (Hs-FN1 310311), COL1A1 (Hs-COL1A1 401891), VCAN (Hs-VCAN 430071), CTSB (Hs-CTSB 490251), COMP (Hs-COMP 457081), COL11A1 (Hs-COL11A1 400741), all from Advanced Cell Diagnostics.

### Isolation and culture of primary cells from human omentum

Fresh tissue was washed in phosphate buffered saline (PBS) and approximately 10cm^3^ of omentum was submerged in 0.25% trypsin (Sigma-Aldrich) and incubated at 37°C for 20min to strip off any mesothelial cells. Trypsin was neutralised using DMEM:F12 1:1 medium (Gibco) with 10% heat-inactivated foetal bovine serum (FBS) (Hyclone). Tissue was washed with PBS, minced with dissection scissors into approximately 1-2mm pieces, suspended in DMEM (Sigma) with 5% FBS and 0.5 mg/ml collagenase type I (Gibco) and placed in a shaking incubator at 50rpm and 37°C for 75min. Tissue digest was passed through 250μM tissue strainers (Thermo-Fisher) and the floating adipocyte layer was carefully collected by pipette and washed by centrifuging twice for 5min at 200*g* in DMEM with 5% FBS. Adipocytes were used immediately for experiments. The stromal vascular fraction (SVF) pellet from the first wash was resuspended in DMEM:F12 1:1 + 10% FBS (growth medium) and cultured at 37°C, 5% CO_2_. After three days, any unattached cells were washed away and attached cells were checked for fibroblastic morphology, and henceforth referred to as omental fibroblasts (OFs). Media was changed every 2-3 days and cells were passaged upon reaching confluence and used for experiments between passages 2-4. Multiple fibroblast donors were used for each experiment and data was plotted for each individual donor with no pooling. In total, fibroblasts from 23 different donors including from tissue with little disease and tissues with confirmed disease.

### HGSOC cell lines

The AOCS1 cell line was established in our laboratory from an omental HGSOC tumor collected during interval debulking surgery in 2011 ^41^. The G164 cell line was established in our laboratory from an omental HGSOC tumor collected during interval debulking surgery in 2015. G164 cells were TP53 and PAX8 positive. Malignant cells were cultured in DMEM:F12 (Gibco), 10% FCS, 1% penicillin and streptomycin in a 5% CO2 humidified incubator at 37°C. The immortalised human FTSE cell line, wild type FT318, was kindly given by Professor Ronny Drapkin (Perelman School of Medicine, University of Pennsylvania) and grown in serum-free WIT-P medium (Cellaria) without antibiotics and 100ng/ml cholera toxin (Sigma-Aldrich). Quality control of all cell lines was carried out by frequent STR analysis (Eurofins MWG), mycoplasma testing (InvivoGen) and cell lines were used for 4 to 5 passages.

### Experimental setups

#### Monocultures

For mRNA extraction and flow cytometry, OFs were seeded at 200k and malignant cells were seeded at 500k in T25 flasks and grown for 4 days. For IF, OFs were seeded at 30k and malignant cells at 60k in 12-wells and grown for 4 days for cell markers and 14 days for matrix molecules. The appropriate factors and inhibitors were added 24h after seeding and replenished every 48h; recombinant TGFβ3 10ng/ml (Peprotech); SB431542 hydrate 20μM (Sigma-Aldrich); GANT61 7.5μM (InSolution, Merck); L-ascorbic acid-2-phopsphate (AA_2_P) 50μg/ml (Sigma-Aldrich). AA2P was only added for experiments involving matrix production.

#### Collagen-gel cultures

Collagen-gel solution (0.1w%) was made for 3D mono- and co-cultures mixing (per 100μl gel) 34μl of 3mg/ml rat-tail collagen I (Gibco), 4μl of 10x DMEM low-glucose (Sigma), 2μl of 1M NaOH and 60μl DMEM:F12 containing cells, prepared on ice. OF-only gels were seeded at 40k and grown for 7 days. Malignant cell-only gels were seeded at 80k and grown for 14 days. Co-cultures were seeded at a ratio of 1:1 (100k:100k) OFs to malignant cells and grown for 7 days. All gels were aliquoted at 100μl in 96-wells and incubated at 37°C, 5% CO_2_for 45min to set and then transferred to free-float in 24-wells with growth medium.

#### Flow-cytometry

For mono-layers, fibroblasts were detached using 0.5% trypsin-EDTA, centrifuged, washed in PBS and resuspended in FACS buffer containing Human Fibroblast Activation Protein alpha PE-conjugated Antibody (FAP-PE) (R&D systems FAB3715P, Clone # 427819) on ice in darkness for 30min. After centrifugation and washing in FACS buffer, cells were suspended in fixation/permeabilization solution (BD Biosciences) for 30 min on ice, washed in permeabilization/wash (PW) buffer, then incubated in 2% goat serum. Human alpha-Smooth Muscle Actin APC-conjugated Antibody (αSMA-APC) (R&D systems IC1420A, Clone #1A4) was added for 30 min before cells were washed in PW-buffer.

For gels containing cells, cultures were digested in 1mg/ml collagenase type I (Thermo-Fisher) in serum-free DMEM for 1hr with shaking at 110 rpm and 37°C. Gels were disaggregated with pipetting and 0.5% trypsin-EDTA (Sigma-Aldrich) was added at 37°C for 30min. DMEM with 10% FBS was added 1:1 to the cell suspension and centrifuged at 200*g* for 5min. For live-dead assay, cells were resuspended in FACS buffer (PBS with 2mM EDTA, 2.5% BSA) containing Fixable Viability Dye eFluor 450 (FVD-e450) (eBioscience 65-0863-18) for 30min on ice protected from light. After washes in FACS buffer, cells were fixed in neutral buffered formalin. Stained samples were analyzed using an LSRFortessa cell analyzer (BD Biosciences) and data were analyzed with FlowJo 9.4.6 (Treestar Inc.).

#### Mechanical characterization of gels

Compression was performed using an Instron ElectroPulse E1000 (Instron, UK) equipped with a 10N load cell (resolution = 0.1 mN). Gels were submerged in PBS throughout testing. Gels were compressed using a stainless steel plane-ended platen with diameter > 2x gel diameter connected directly to the load cell. Gel thickness was measured as the distance between the base of the test dish and top of the gel, each detected by applying a pre-load of 0.3-5 mN. Tests were performed in displacement control mode and gels were displaced to 30% thickness at a rate of 1%s^-1^ with the resulting load recorded. Gel modulus, a measure of material stiffness independent of specimen geometry, was calculated by converting load-data to stress (kPa) (load ÷ gel area), plotting a stress-strain curve and then taking the slope of the curve between 15-20% strain.

#### RNA isolation and real-time quantitative PCR

Total RNA was extracted using Qiagen RNeasy Plus Micro kit according to the manufacturer’s instructions. Monolayers were first scrapped in RLT Plus buffer (Qiagen) and RNA was quantified using a NanoDrop 2000c (Thermo-Fisher Scientific). Tri-cultures were placed directly into RLT buffer and rigorously vortexed. RNA quality was analyzed on Agilent bioanalyzer 2100 using RNA PicoChips according to manufacturer’s instructions. RNA integrity numbers were between 8.1 and 9.9. Total and reverse-transcription was carried out on 1μg of RNA using a T100 Thermal Cycler (Bio-Rad) and a High-Capacity cDNA Reverse Transcription Kit (Applied Biosystems) according to manufacturer’s instructions. The PCR reaction was run on a StepOnePlus Real-Time PCR System (Applied Biosystems) using iTag Universal Probes Supermix (Applied Biosystems), FAM-MGB labelled Taqman gene expression probes and 5ng sample cDNA. Taqman gene expression assay targets; ACTA2 (Hs00426835_g1), FAP (Hs00990791_m1), GLI1 (Hs00171790_m1), GLI2 (Hs01119974_m1), GLI3 (Hs00609233_m1), TGFβ1 (Hs00998133_m1), TGFβ2 (Hs00234244_m1), TGFβ3 (Hs01086000_m1), FN1 (Hs01549976_m1), COL1A1 (Hs00164004_m1), VCAN (Hs00171642_m1), CTSB (Hs00947433_m1), COMP (Hs00164359_m1), COL11A1 (Hs01097664_m1); GAPDH (Hs027866254_g1) and 18S (Hs03003631_g1), were both used as housekeeping genes (All from Thermo-Fisher Scientific, UK).

#### Preparation of adipocyte-collagen gels

Purified adipocytes (1ml) were combined with the following reagents to give 0.1w% collagen-adipocyte gels: 1ml of 3mg/ml rat-tail collagen I, 100μl of 10x DMEM low-glucose, 48μl of 1M NaOH and 852μl H_2_O, prepared on ice. Adipocyte-gel mixture was incubated at 37°C, 5% CO_2_for 45min in 100μl aliquots in a 96-well dish. Gels were gently transferred to 24-wells and cultured in 1ml Medium-199 with insulin-transferrin-selenium (Gibco). Adipocyte gels were used for experiments with other cell types within 7 days of isolation. For live/dead assays, gels were immersed in 1ml PBS containing 20μg fluorescein diacetate (Sigma-Aldrich) and 4μM ethidium homodimer (Sigma-Aldrich), incubated for 10min at 37°C, then placed on a glass slide with PBS to prevent drying. Gels were imaged using a Zeiss LSM510 confocal microscope.

#### Preparation of tri-cultures

3D co-cultures of fibroblasts and HGSOC cells were first cultured for 7 days to allow gel contraction and remodelling. Adipocyte gels were placed in dry 24-wells and then using a Pasteur pipette, a co-culture gel with a small volume of media was placed on top of the middle of an adipocyte gel. Co-cultures were embedded into the centre of adipocyte gels by carefully pressing down with the curved end of a sterile 1.5ml eppendorf. Wells were then filled with culture media and tri-cultures were allowed to free-float. After 24 hours, ascorbic acid (50μg/ml) was added to gels without or with inhibitors, SB431542 (20μM) and GANT61 (7.5μM). Gel images were acquired before adding factors and at the end of culture (14 days). Media and factors were replenished every 2-3 days and cultured for a further 13 days. At the end point (21 days total culture), tri-cultures were washed well with PBS and fixed in 10% formalin for 2h for IF, or for 24h for paraffin embedding. After fixation, gels were stored in PBS at 4°C until processed.

#### Immunohistochemistry

FFPE sections (4 μm) of omentum samples or gel-cultures were re-hydrated in ethanol solutions: 100%, 90%, 70%, and finally 50%. Sections were transferred to citric acid-based antigen unmasking solution (Vector Laboratories) and heated in a 2100 antigen-retriever (Aptum Biologics). Sections were treated with 3% H2O2 for 5min and blocked with 5% BSA for 1hr. Primary antibody was added in antibody diluent (Zytomed) for 1hr. Slides were washed and a biotinylated secondary antibody (Vector) was added. Subsequent steps were carried out according to the protocol included with the Vectastain Elite ABC HRP kit. Slides were incubated for 5min with DAB solution made using Sigmafast DAB tablets (Sigma-Aldrich). Finally, slides were counterstained in 50% Gill’s haematoxylin I, and dehydrated in 50%, 70%, and 100% ethanol then twice in xylene. Coverslips were affixed using DPX mountant (Sigma-Aldrich). All sections were scanned using a 3DHISTECH Panoramic 250 digital slide scanner (3DHISTECH), and the resulting scans were analyzed using Definiens software (Definiens AG). Disease scores were determined first by manually defining regions of interest in the tissue that represented tumor, stroma, fat (adipocytes), and then training the software to recognize these regions of interest. Disease score was expressed as a percentage of the whole tissue area that contained tumor and/or stroma (Fig. 1A).

#### Immunofluorescence

Gels were fixed overnight in 10% neutral buffered formalin, washed in PBS and permeabilized in Triton X-100 (0.5% in PBS) for 10min. Gels were incubated in blocking solution (5% BSA or goat serum), and then incubated with primary antibody overnight at 4°C. Gels were washed and incubated with fluorescent secondary antibody, for 1hr protected from light and then washed. Finally, gels were incubated with 0.4 μg/ml DAPI and then washed. Fluorescent images of tri-cultures were captured on an inverted Zeiss LSM 510 laser-scanning confocal microscope using a 10x or 20x air objective. Specimen images were acquired with a field of view equal to 238.1 × 238.1μm containing 1024×1024 pixels. All imaging conditions including laser settings and scan settings were kept constant for all gel groups for each fluorescent-labelled antibody. Images of monolayers were captured using an EVOS FLoid Cell Imaging Station. For F-actin, Alexa Fluor 568 Phalloidin (A12380, Thermo Fisher Scientific) was used.

#### Antibodies

The following antibodies were used for immunostaining: anti-actin, α-smooth muscle (clone 1A4, A2547), anti-VCAN (polyclonal, HPA004726), anti-COL11A1 (polyclonal, HPA052246), anti-COL1A1 (polyclonal, HPA011795), anti-FN1 (polyclonal, F3648) all from Sigma-Aldrich, UK; anti-Ki67 (cloneMIB-1, M7240), from Dako, UK; anti-fibroblast activation protein, alpha (EPR 20021, ab207178), anti-COMP (ab11056), anti-CTSB (CA10, ab58802), anti-TGFβ all from Abcam; anti-PAX8 (NBP1-32440) from Novus; anti-EPCAM Alexa Fluor 488 conjugated (53-8326-41) from Thermo-Fisher; GLI-1 Antibody (C-1, sc-515751) from Santa Cruz Biotechnology.

### Statistical and Bioinformatics Analysis

Statistical analyses and graphics were performed in GraphPad Prism or the programming language R (version 3.1.3). All correlations were calculated using Spearman’s rank correlation. For pairwise comparisons a two-way paired t-test was used. For comparisons of >t2 sample means, one-way ANOVA with Tukey’s HSD test were used. Differential expression analysis was performed in Edge R using limma (PMID:25605792). Gene-set enrichment analysis (GSEA) was performed using the GSEA software ^42^ to identify the canonical pathways gene sets from the Molecular Signatures Database (MSigDB-C2-CP v6.2). See figure legends for significance levels and number of samples, n. For experiments involving OFs, n represents technical replicates and N represents number of donors. Data were considered statistically significant from *p* < 0.05.

#### ICGC analysis for six matrisome molecules

The ICGC_OV read counts across 93 primary tumors were extracted from the exp_seq.OV-AU.tsv.gz file in the ICGC data repository Release 20 (http://dcc.icgc.org). Only genes that achieved at least one read count in at least ten samples were selected, producing 18,010 filtered genes in total. Variance stabilizing transformation was then applied using the rlog function ^43^. Overall survival (OS) was extracted from the donor.OV-AU.tsv.gz file. Mean expression for the six matrisome genes was calculated for each sample and high and low matrix groups were determined using the method described previously ^44^. Survival modeling and Kaplan-Meier (KM) analysis was undertaken using R package survival. OS was defined as time from diagnosis to death, or to the last follow-up date for survivors. The significantly differentially expressed genes were selected using a false discovery rate (FDR) < 0.05.

#### RNA-seq and analysis

RNA-seq was performed by the Wellcome Trust Centre for Human Genetics (Oxford, UK) to approximately 30x mean depth for the HGSOC cell lines or 20x for the 3D cultures. The sequencing was carried out on the Illumina HiSeq4000 or on the NovaSeq6000 platform, strand-specific, generating 150bp paired-end reads. RNA-Seq reads were mapped to the human genome (hg19, Genome Reference Consortium GRCh37) in strand-specific mode as part of the Wellcome Trust Centre pipeline. Number of reads aligned to the exonic region of each gene were counted using htseq-count based on the Ensembl annotation. Only genes that achieved at least one read count per million reads (cpm) in at least twenty-five percent of the samples were kept. Conditional quantile normalisation was performed counting for gene length and GC content and a log_2_transformed RPKM expression matrix was generated. RNA-Seq data have been deposited in Gene Expression Omnibus (GEO) under the accession number GSE125109.

## Acknowledgements

The authors wish to acknowledge the role of the Barts Cancer Institute Tissue Bank Team in collecting and making available the samples used for this publication. We thank Barts Trust Oncology Surgeons for sample provision. We also thank Dr Chiara Berlato, Anissa Lakhani, Joash Joy and Priyanka Hirani for technical help, George Elia and the BCI Pathology Core, Professor David Bowtell and Professor Ronny Drapkin for kindly providing cell lines. Finally, we express our gratitude to the patients for donating the samples without which this work would not have been possible. This project was funded by the European Research Council (ERC322566) and Cancer Research UK (A16354, A13034, A19694 and A25714). Laura Lecker was funded by a grant from the Institute of Bioengineering, Queen Mary University of London.

