## Supplementary material for "Modelling TGFβR and Hh pathway regulation of prognostic matrisome molecules in ovarian cancer": Suppl Figures

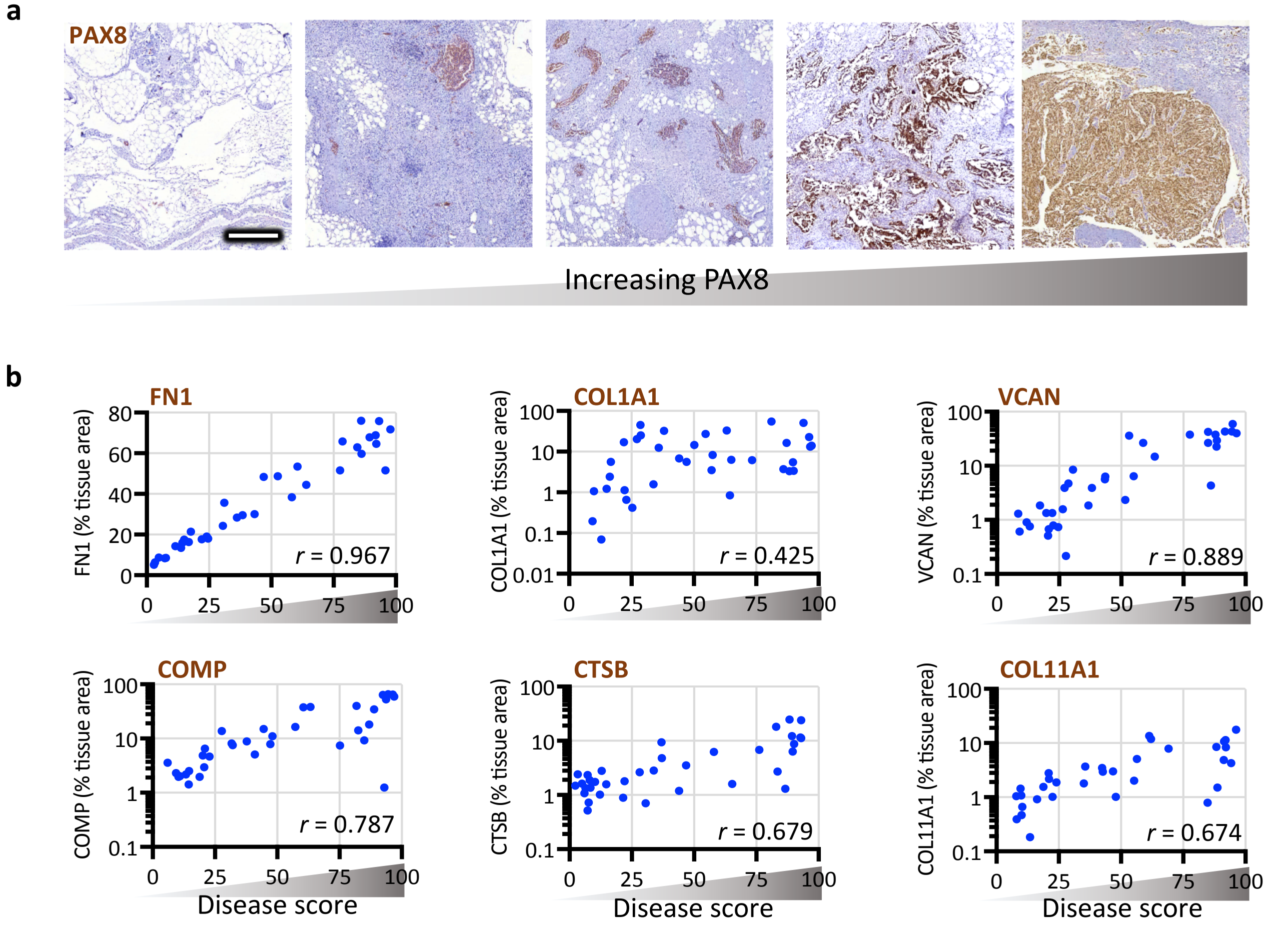


**Supplementary Figure 1. HGSOC tissue biopsies were PAX8, TGFβ, αSMA and FAP positive.** (**a**) HGSOC tissue biopsies had varying levels of PAX8 positivity and stromal remodelling. (**b**) Area of tissue IHC stain for FN1, COL1A1, VCAN, COMP, CTSB, COL11A1 was quantified on HGSOC biopses; all showed positive correlations with disease score (Spearman's rank correlation coefficient, *r*). Scale bars are 500µm.

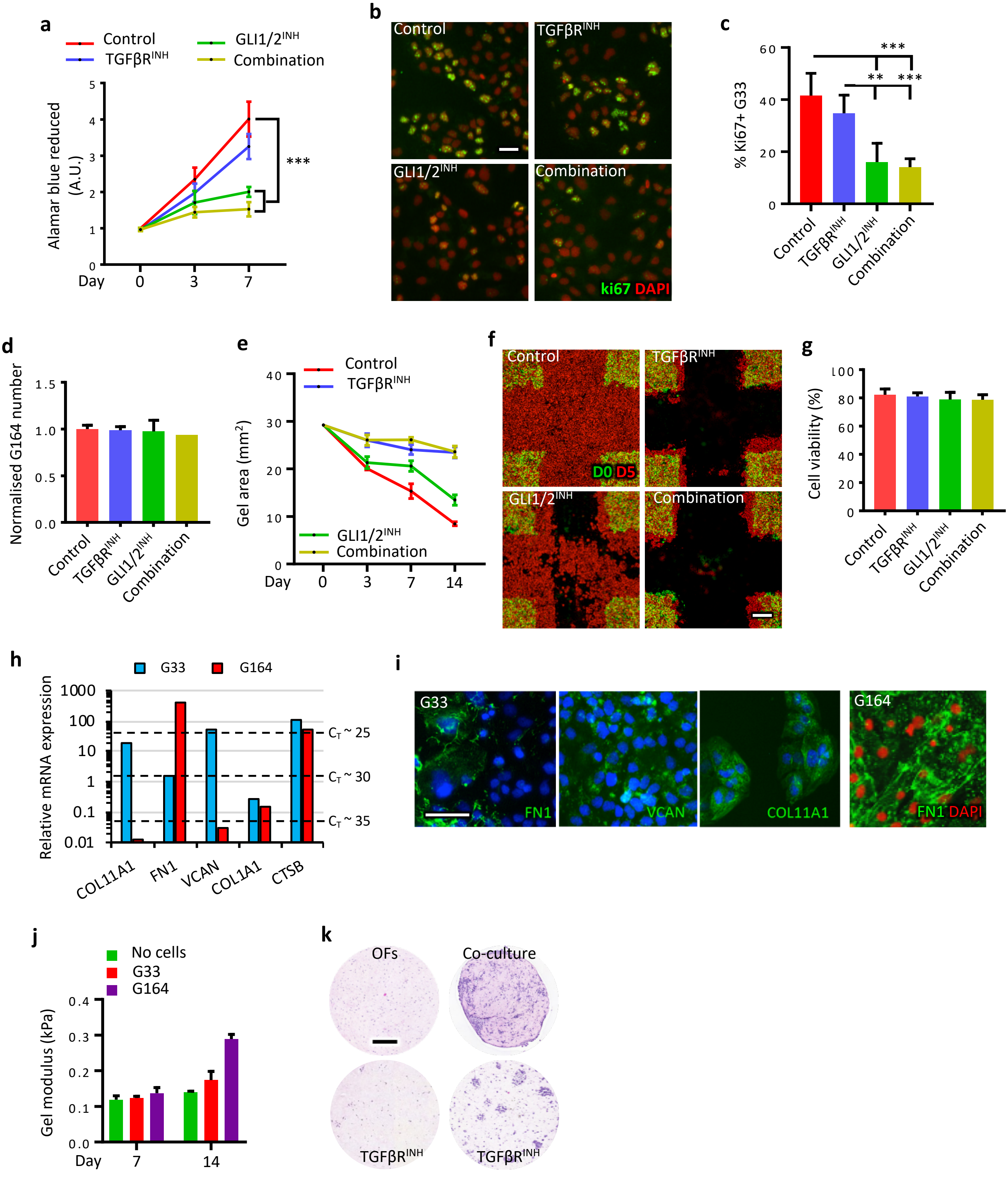


**Supplementary Figure 2. Hh and TGFβR pathway activity as well as matrix molecule expression is malignant cell dependant.** (**a-c**) G33 were cultured for 7 days with SB431542 (TGFβR$^{INH}$), GANT61 (GLI1/2$^{INH}$) or combination. (**a**) Alamar blue assay was performed at 2h (day 0), days 3 and 7; data is mean ± SD (N = 3), p < 0.001 (one-way ANOVA-Tukey's). (**b**) Ki67 IF of G33 at 3 days and (**c**) cells with nuclear positivity were counted (data is mean ± SEM for n>200 cells, N = 3, **p < 0.01, ***p < 0.001 (one-way ANOVA-Tukey's). (**d-f**) G164s grown in COL1 gels for 14 days alone or with TGFβR$^{INH}$, GLI1/2$^{INH}$ or combination. (**d**) G164 number assessed via flow cytometry (mean ± SD, N = 2) and (**e**) gel area measurements (mean ± SD, n=4 N=2) and (**f**) cell viability. (**g**) A scratch assay was performed on high density G164s and images were acquired directly after (D0) and 5 days (D5) later; images were pseudo coloured and overlayed. (**h**) Matrix molecule expression analysed via qRTPCR for G33 and G164; dashed lines and $C_T$ number represent qRTPCR cycle number. (**i**) IF staining performed on G33 for FN1, VCAN and COL11A1 and on G164 for FN1. (**j**) Modulus of COL1 gels at days 7 and 14, containing no cells or malignant cells only (mean ± SD, n=3, N=3). (**k**) OFs were cultured in 3D COL1 gels alone or as co-cultures and grown for 7 days with or without TGFβR$^{INH}$and H&E was performed on fixed sections. Scale bars are (**b, i**) 50µm, (**g**) 100µm and (**k**) 250µm.

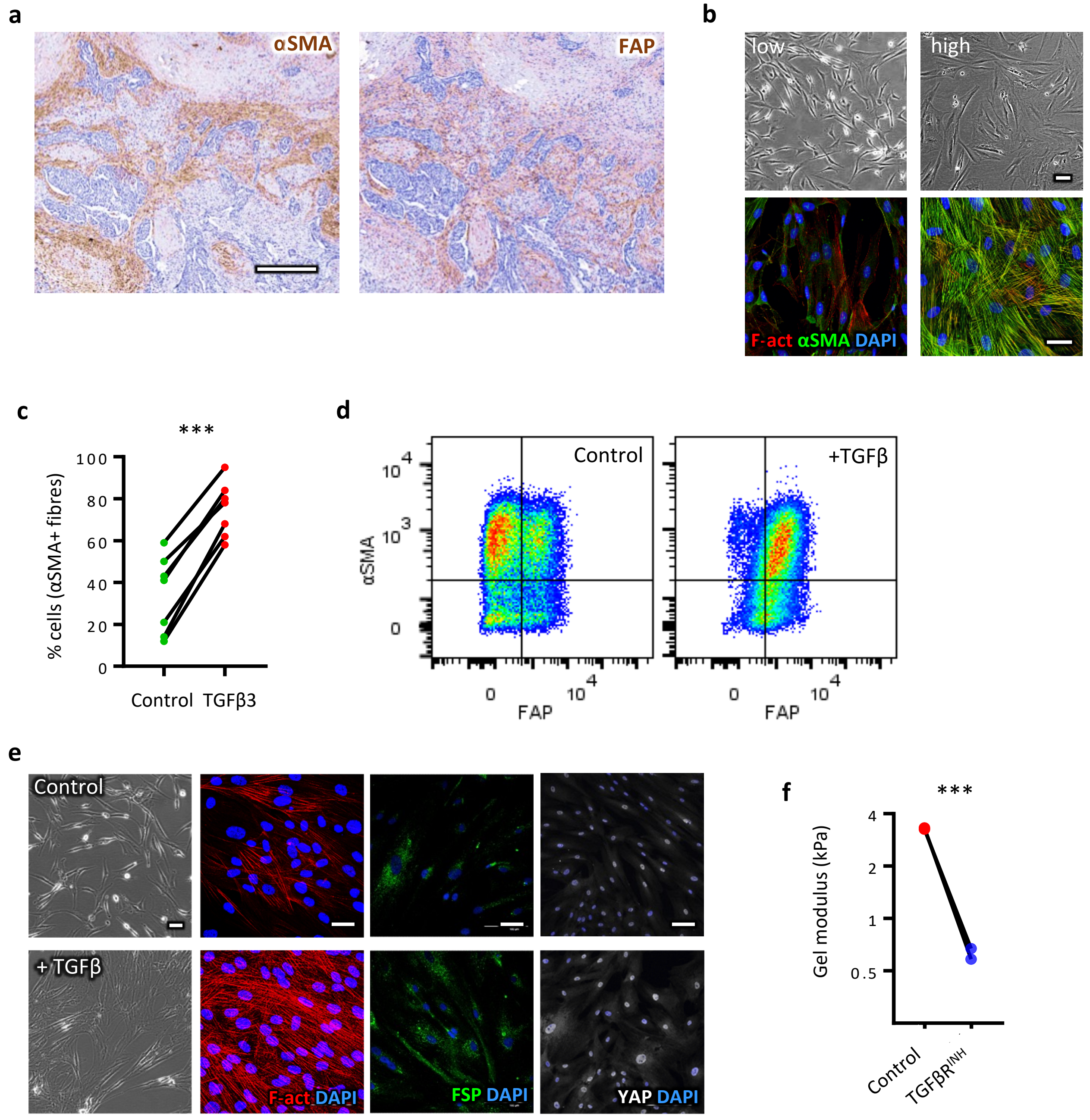


**Supplementary Figure 3. TGFβ3-treated omental fibroblasts display phenotypic changes.** (**a**) Sequential sections were taken from a biopsy with high disease score and stained via IHC for αSMA and FAP. (**b**) Omental fibroblasts (OFs) were isolated from HGSOC patients and described by low (L-OFs) or high (H-OFs) activation based on morphology and level of expression of F-actin and αSMA fibres observed via phase contrast and IF, respectively. (**c**) A significant increase in the proportion of OFs with αSMA fibers treated with TGFβ3 was measured versus controls across 7 donors (n>100 cells per donor), $p < 0.001$ (two-way, paired t test). (**d**) Flow cytometry performed on control and TGFβ3-treated OFs stained for αSMA and FAP; dot plot displays a representative example of collected and sorted viable cells gated for both markers. (**e**) OFs treated with TGFβ3 appeared larger, more spread, and show increased expression of F-actin, FSP (fibroblast surface protein) and YAP via IF staining compared with L-OFs (control). (**f**) Gels containing H-OFs had a significantly decreased modulus with SB431542 ($TGFβR^{INH}$) treatment (mean, N=2 (n=4), $p < 0.001$ (two-way, paired t test). Scale bars are (**a**) 200µm and (**b, e**) 50µm.

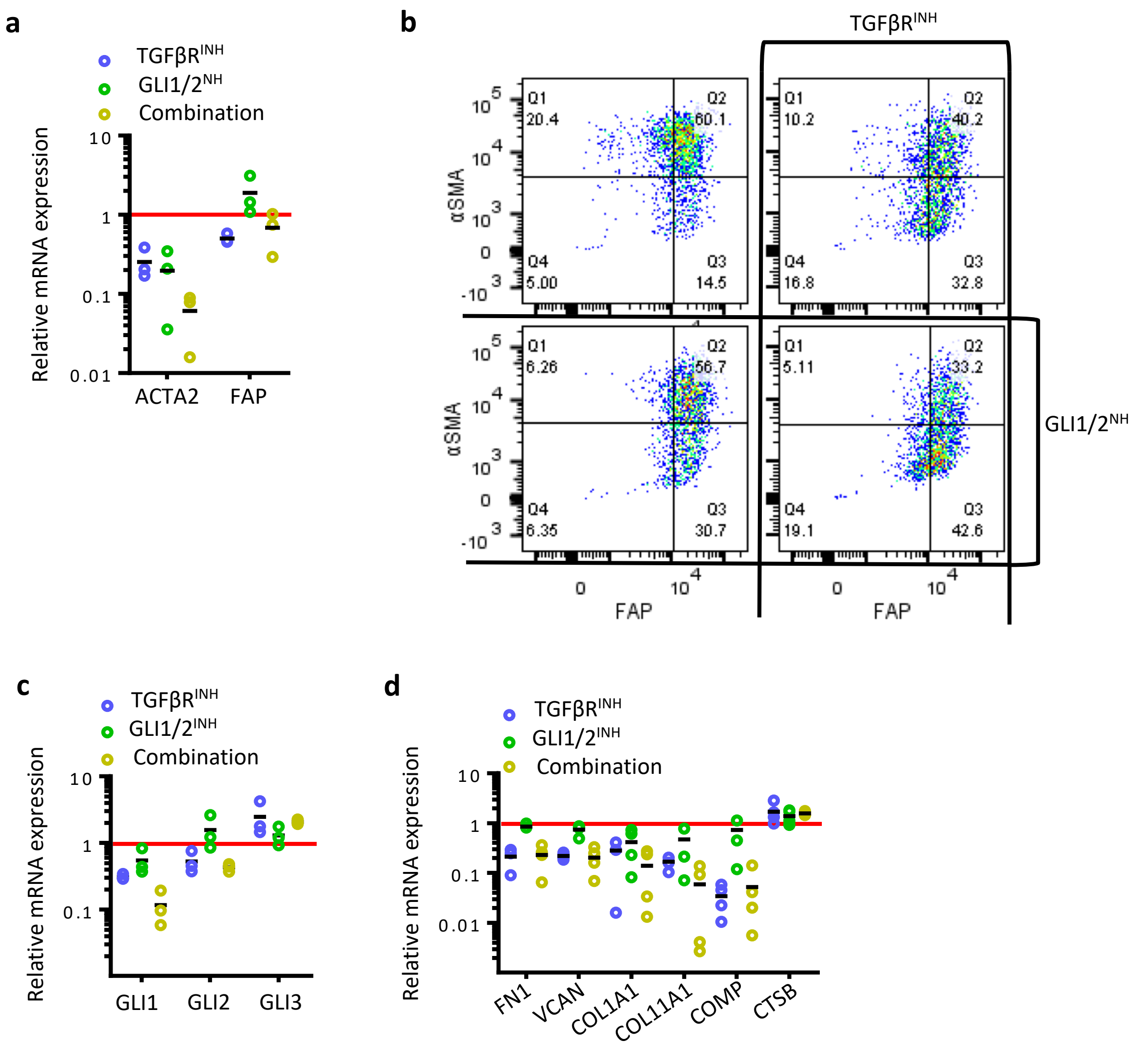


**Supplementary Figure 4. TGFβRI and GLI1/2 inhibitors reduce omental fibroblast activation and matrix expression.** (**a-d**) L-OFs treated with TGFβ3 and cultured with SB431542 (TGFβRI$^{INH}$) or GANT61 (GLI1/2$^{INH}$) inhibitors separately or in combination; (**a**) qRT-PCR was performed to assess ACTA2 and FAP expression (N = 3); (**b**) Flow cytometry was performed to assess αSMA and FAP positivity on viable cells (top left is control then clockwise TGFRI$^{INH}$, GLI1/2$^{INH}$, combination); (**c**) qRT-PCR performed to assess GLI1/2/3 expression (N = 3) and (**d**) matrix molecule expression (N = 4). Red lines represent control (no treatment) expression. Scale bar is 50µm.

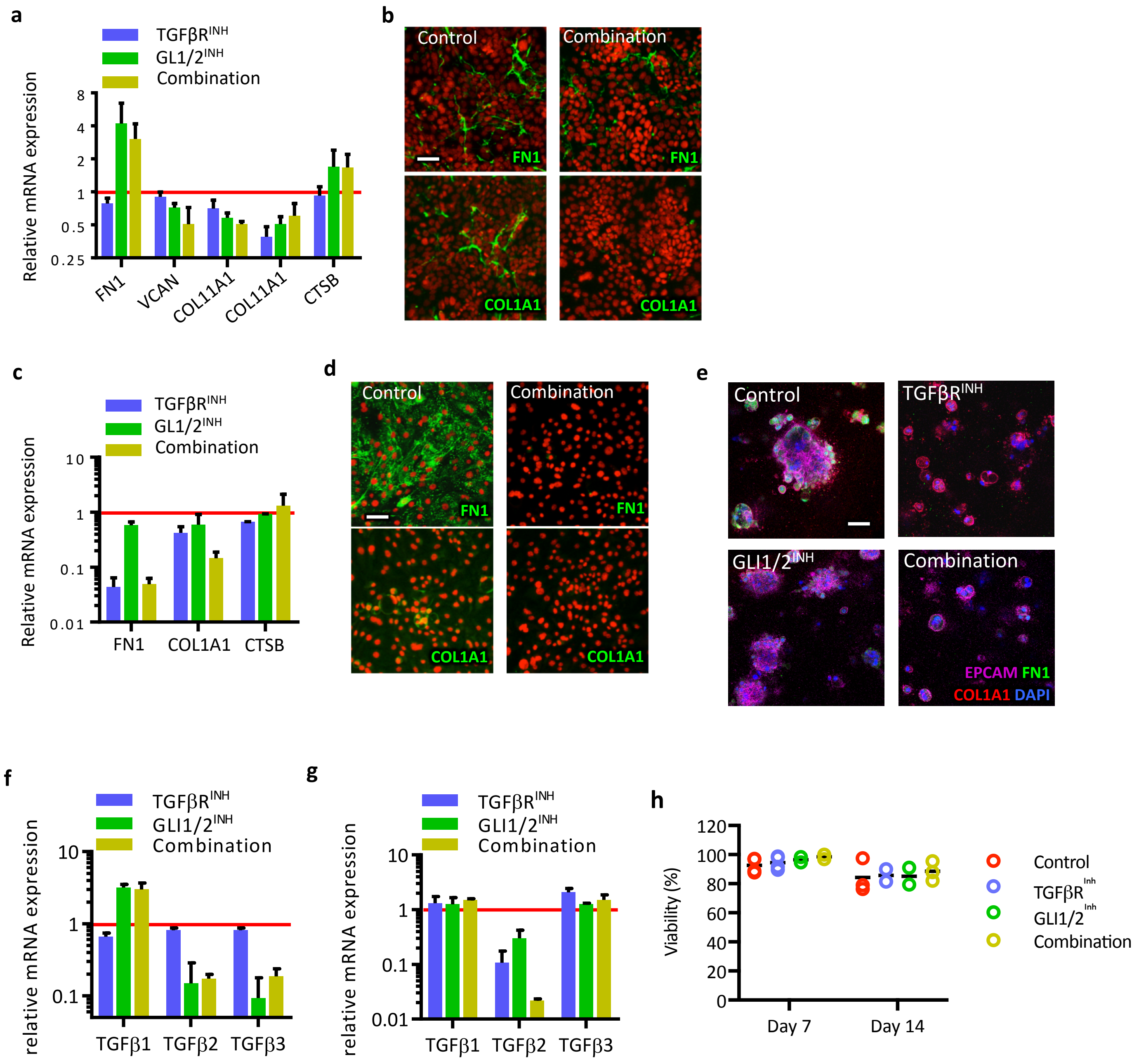


**Supplementary Figure 5. HGSOC malignant cells heterogeneously express matrix molecules.** (**a-b**) G33 and (**c-e**) G164 cells were treated with SB431542 (TGFβR$^{INH}$) or GANT61 (GLI1/2$^{INH}$), or combination. (**a**) Matrix molecule expression analysed via qRTPCR in G33;. (**b**) IF staining for FN1 and COL1A1 performed on G33 with/without combination treatment. (**c**) Matrix molecule expression analysed via qRTPCR in G164s. (**d**) IF staining for FN1 and COL1A1 performed on G164 with/without combination treatment. (**e**) G164s were cultured in 3D COL1 gels for 14 days and stained for EPCAM, FN1, and COL1A1 via IF. (**f**) TGFβ1/2/3-isoform mRNA expression for G33. (**g**) TGFβ1/2/3-isoform mRNA expression for G164. (**h**) Cell viability of 3D cocultures were assessed with a fixable cell viability dye by flow cytometry at days 7 and 14 for control gels and gels with inhibitors (n = 2-3). All data is mean ± SD (n = 3) and normalised to no-treatment control (red line). Scale bars are 50μm.

a

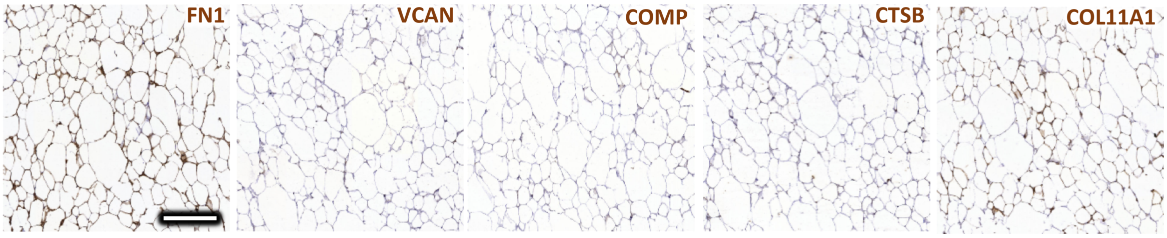


b

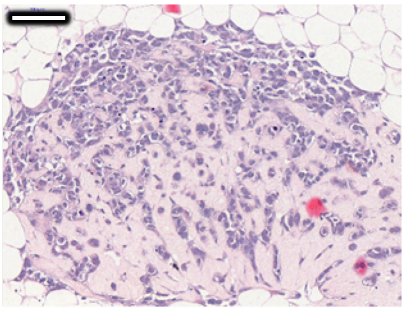

**Supplementary Figure 6. Adipocyte gels display low levels of tumour-matrix proteins.** (**a**) Adipocyte gels fixed and stained via IHC for matrix proteins FN1, VCAN, COMP, CTSB, and COL11A1. (**b**) H&E of Tri-culture model fixed-section after 21 days of culture shows tumour core. Scale bar is (**a**) 200µm and (**b**) 100µm.

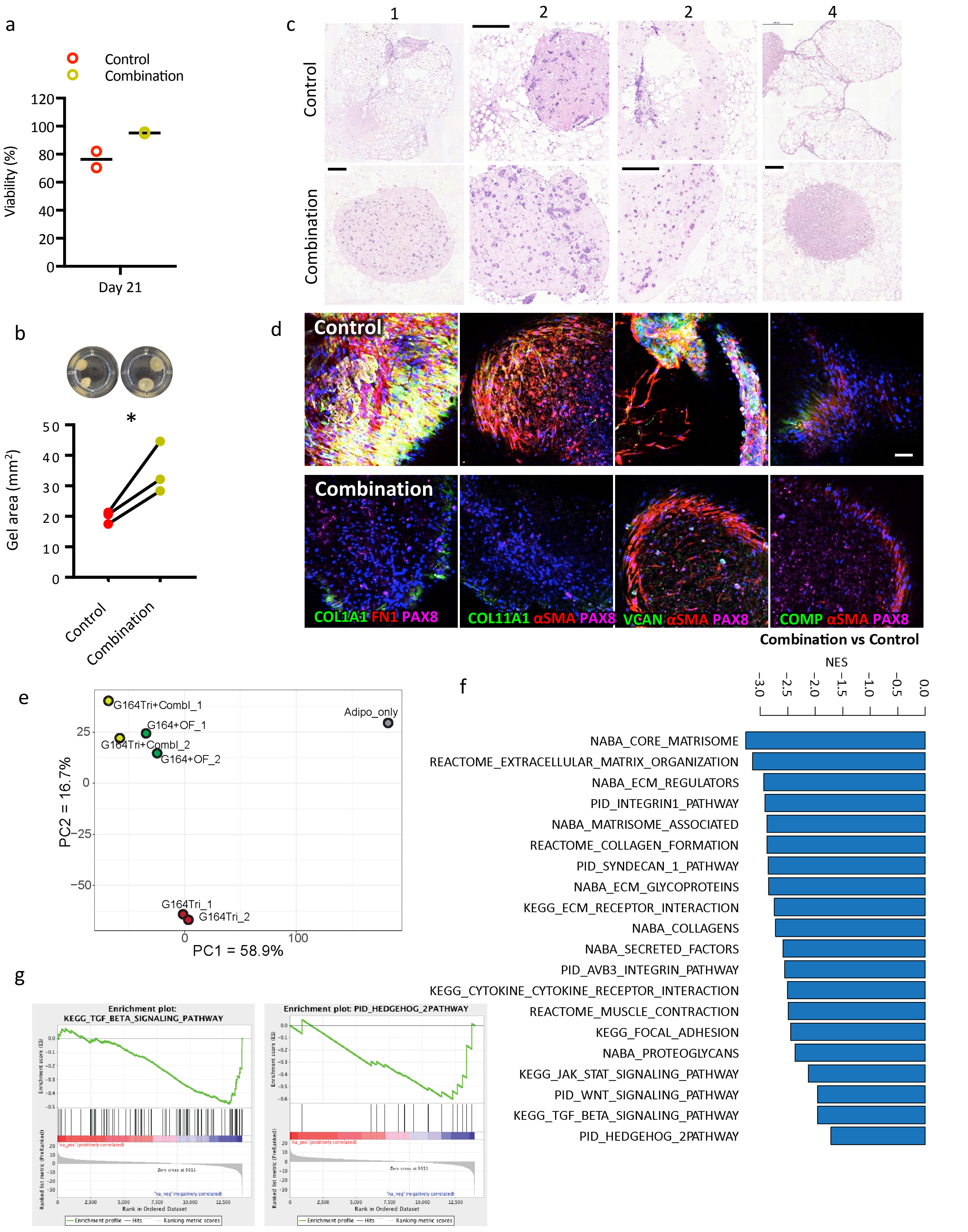


**Supplementary Figure 7. TGFβRI and GLI1/2 inhibitor combination reduce tri-culture remodelling.** (a) Viability assessed at day 21 for control and combination-treated tri-cultures using viability dye via flow cytometry and (n = 2). (**b**) Gel area of G33 tri-cultures measured at day 21 (mean for 3-5 gels per experiment, n = 3; two-way, paired t test, $p < 0.05$). (**c**) Tri-cultures with or without combination-treatment were fixed at 21 days, sectioned and stained with H&E; images show four different experimental examples, 1-3: G164 tri-cultures and 4: G33 tri-culture. (**d**) G33 tri-cultures IF stained for matrix molecules FN1, VCAN, COL1A1, COMP and COL11A1, and αSMA at 21 days of culture. (**e**) Principal component analysis for RNASeq of tri-cultures. (**f**) Normalised enrichment scores derived from GSEA of RNAseq data of G164Tri+CombI vs G164Tri ($p < 0.05$). (g) Enrichment plots derived from GSEA of RNAseq data of G164 tri-cultures. 2 gels were combined per sequencing sample, Adipo_only (n = 1), G164Tri and G164Tri+CombI (n = 2). Scale bars for (**b**) are 100μm and (**d**) are 250μm.
